# Mechanical conditions preventing live cell extrusion during primary neurulation in amniotes

**DOI:** 10.1101/2025.08.08.668972

**Authors:** Santiago A. Bosch-Roascio, Julio A. Hernández, Flavio R. Zolessi

## Abstract

During primary neurulation in amniote embryos, the neural plate gives rise to the neural tube in a process requiring the coordination of forces at different scales throughout a geometrically complex tissue. The ways in which this process fails inform us of the complex mechanical conditions required for its correct completion. Previous results showed that the functional disruption of MARCKS, a protein which simultaneously interacts with the plasma membrane and actin filaments, resulted in neural tube closure defects with apical cell extrusion. Here, we demonstrate that this is an example of “live cell extrusion”, wherein extruded cells are not undergoing apoptosis. This suggests that extrusion in this case might be due to a mechanical instability in the neural plate. Using an expanded energy-based vertex model of pseudostratified epithelia we then show that extrusion may be elicited by a reduction in the relative surface tension of apical and basal interfaces with respect to cell-cell interfaces. Finally, by considering a continuum description of a simplified epithelium we derive an approximate quantitative threshold for single-layered epithelial stability in the form of a power law relating cell density to the relative value of interfacial surface tensions. Our work serves to explain an example of how alterations in polarization and forces at the single-cell level can produce tissue-scale instabilities which not only greatly alter its morphology but can also ultimately lead to severe developmental defects.

**Highlights:**

- PMA treatment induces live apical cell extrusion in the chick embryo neural plate
- Our energy-based vertex model mechanically simulates a pseudostratified epithelium
- Higher apical/basal compared to lateral surface tension leads to epithelial stability
- Lower apical/basal surface tension leads to epithelial instability and cell extrusion
- A power law between cell density and surface tension defines the stability threshold

## 1. Introduction

In vertebrates, the central nervous system almost entirely derives from a specialization at the dorsal ectoderm, termed the neural plate, an epithelium that eventually deforms to ingress into the embryo as a whole and forms the neural tube. Particularly in amniotes, this “primary neurulation” process involves the invagination and folding of the neuroepithelium along the cephalo-caudal axis of the embryo to become a tube separated from the epidermal ectoderm that remains at the surface. The series of morphogenetic movements that drive neurulation arise from intrinsic and extrinsic forces exerted over the neuroepithelial cells, which all along the process remain cohesive and highly polarized (Colas and Schoenwolf, 2001; de Goederen et al., 2022). The intrinsic forces deforming the neural plate are generated at different levels, including actions mediated by the cytoskeleton. In addition, forces extended along several cells in the neural plate, such as those generated by convergent-extension movements (Schoenwolf and Smith, 1990), or coming from the surrounding non-neural ectoderm (Alvarez and Schoenwolf, 1992; Moury and Schoenwolf, 1995), could contribute to bending. In this regard, the early work by Lee and Nagele has indicated that neural plate-intrinsic forces would be sufficient for the formation of the neural tube in chick embryo explants (Lee and Nagele, 1988).

All these forces are applied to cells that, in addition, greatly increase their proliferation rate from the neural plate stage on, becoming a quickly growing population (Copp and Greene, 2010). Cells remain attached to both apical and basal surfaces, but they increase their height and reduce their width, generating a tall pseudostratified epithelium. In this organization, cells display interkinetic nuclear migration, by which cell divisions occur at the apical border of the neuroepithelium, while S-phase tends to occur in nuclei closer to the basal side (Norden, 2017; Sauer, 1936). The combination of all these factors results in neuroepithelial cells with extremely complex shapes (Gómez et al., 2021; Iber and Vetter, 2022).

How is it possible that this epithelial cell population tolerates the enormous forces that drive neurulation while proliferating at a high rate, without disruption? Different lines of evidence indicate that the key might be in the strict maintenance of apico-basal cell polarity. Among other factors, this is achieved through a tight binding of each cell to the basal lamina on the basal side, and to their neighboring cells at the sub-apical border through integrin- and cadherin-based cell adhesions, respectively (Saade and Martí, 2025). These adhesion complexes tightly, but dynamically, bind to the cytoskeleton, establishing a mechanically continuous meshwork that spans across cells, and that can be modulated in response to different external and internal cues (Campàs et al., 2024; Sluysmans et al., 2017; Zhang and Wei, 2025). It has been generally accepted that microtubules extend during neurulation, elongating cells along the apico-basal axis, while the actin cytoskeleton acts in different ways, allowing cells to take a wedge-like shape (Cearns et al., 2016; Schoenwolf and Powers, 1987). The maximal accumulation of actin filaments is observed close to the apical cell border, in association with the cadherin-based *zonula adherens*, and acto-myosin contraction at this level has been indicated as an important motor in apical surface reduction (Ampartzidis et al., 2023; Escuin et al., 2015; Martin and Goldstein, 2014; Sawyer et al., 2011). This apical constriction would be more relevant in highly-bent areas of the neural plate, when they exist, such as the ventromedial and the dorso-lateral hinges. Here, other cell processes, like basal positioning of nuclei and localized programmed cell death were also shown to contribute to epithelial bending, highlighting the importance of regulating cell crowding during neurulation (Roellig et al., 2022; Smith and Schoenwolf, 1988).

It is not surprising, then, that several polarized actin-modulating proteins have been demonstrated to be necessary for primary neurulation. Among these are the MARCKS family members. MARCKS (Myristoylated Alanine Rich C-Kinase Substrate) and MARCKS-Like 1 are ubiquitous proteins present only in vertebrates, which simultaneously interact with the plasma membrane and actin filaments (Arbuzova et al., 2002; El Amri et al., 2018). Phosphorylation by PKC (or interaction with calcium-calmodulin) causes MARCKS to detach from the membrane and to lose one of the two actin-binding sites (McLaughlin and Aderem, 1995). Hence, MARCKS acts as a molecular switch connecting extracellular signals to the cell cortex. Disruption of MARCKS family protein expression causes severe neurulation defects in both mice and zebrafish embryos (Chen et al., 1996; Prieto and Zolessi, 2017; Stumpo et al., 1995). Further, we found that MARCKS is accumulated at the apical membrane in neural plate cells of the chick embryo only during neural tube closure, and that its knockdown causes a failure of neural tube closure (Aparicio et al., 2018; Zolessi and Arruti, 2001). Interestingly, the local cellular phenotype observed in these embryos was very similar to the effect obtained after treatment with a strong agonist of PKC, the phorbol ester PMA: loss of apico-basal cell polarity accompanied by apical cell extrusion. This effect of PMA was effectively rescued by the expression of a non-phosphorylatable form of MARCKS, where all serines at the ED were substituted by asparagines (Aparicio et al., 2018).

Apical cell extrusion, that is, the expulsion of a cell through the apical side of the epithelium, has been generally reported as an active mechanism for maintaining epithelial density and integrity, and involves biochemical and mechanical signaling such as the secretion of sphingosine-1-phosphate (S1P) by extruding cells, which activates Rho and actomyosin in neighboring cells (Eisenhoffer and Rosenblatt, 2013; Nanavati et al., 2020). One function of this process is the elimination of dying cells from the epithelium (“apoptotic cell extrusion”), and implies medium-scale epithelial topological irregularities, pulsatile contractions and loss of tissue tension (Atieh et al., 2021; Saw et al., 2017), together with the local activation of lamellipodial protrusions and the assembly of a basally-localized contractile actomyosin ring, in a cadherin-dependent manner (Duszyc et al., 2021; Lubkov and Bar-Sagi, 2014; Rosenblatt et al., 2001). Conversely, “live apical cell extrusion” appears as a counterbalancing mechanism to avoid cell overcrowding in a proliferating cell population, and occurs regularly in many physiological processes of epithelia, like in cell shedding in intestinal microvilli and in the control of carcinogenic transformation (Madara, 1990; Tanimura and Fujita, 2020). It requires stretch-activated ion channels (Piezo), S1P and the Rho-myosin pathway, and also produces regular topological changes in the organization of the epithelium, based on the collective action of surrounding cells (Eisenhoffer et al., 2012). During this type of extrusion, the cell being expelled remains alive and plays an active part in the process, eventually dying by anoikis after the extrusion has been completed (Marinari et al., 2012; Slattum and Rosenblatt, 2014).

Taking all this information, we wondered if apical cell extrusion of neuroepithelial cells upon MARCKS knockdown or PKC activation could be a live cell extrusion mechanism related to an imbalance in the mechanical forces in the cell population. This could be caused either by the alteration of the actin cytoskeleton or by the downregulation of apico-basal cell polarity. By pharmacologically inhibiting caspase activity on PMA-treated chick neurulas, we demonstrate here that the observed massive apical cell extrusion is mediated by a live mechanism. Secondly, we aimed at explaining the experimental observations by generating a theoretical model and performing computational simulations. We present an expanded 2D vertex model of a pseudostratified epithelium, to show that alterations in the balance of forces imposed to cells on the apical or basal plasma membrane domains compared to lateral domains, was enough to robustly obtain situations that could only be solved by massive cell extrusion.

## 2. Methods

### 2.1. Embryo culture

Wildtype chicken eggs were provided by a local producer (PRODHIN) and incubated at 37° C for at least 28 h, corresponding to stage HH 8 (Hamburger and Hamilton, 1951). Chick embryos were cultivated *ex ovo* with ventral side up over a semi-solid agar/albumen substrate in 35 mm Petri dishes using the EC culture technique (Chapman et al., 2001). Cultured embryos corresponding to stages lower than HH8-/8 were incubated until they reached the desired stage. A total of 72 embryos were processed, spread across five experiments.

### 2.2. Pharmacological treatments

Cultured embryos were divided into four groups according to treatment: those receiving treatment with PMA, QVD-OPh, neither (control) or both (double, PMA+QVD). Stock solutions were prepared for PMA (Sigma-Aldrich P-8139) at 5 mM and QVD-OPh (MedChem Express #HY-12305) at 20 mM, both diluted in 100% DMSO.

All embryos were pre-incubated for 30 min. at 37° C with a 20 μl drop containing either 0,83

% DMSO in 1x PBS (control and PMA) or 167 μM QVD-OPh (1:120 dilution; QVD-OPh and double), applied ventrally. Afterwards, the excess liquid was removed and embryos were moved to a new culture dish containing 0.93% DMSO (control), 5 μM PMA (1:1000 dilution) and 0.83% DMSO (PMA), 167 μM QVD-OPh and 0.1% DMSO (QVD-OPh) or PMA and QVD-OPh (double) embedded in the substrate. Additionally, a new 20 μl droplet containing either DMSO or QVD-OPh was applied as before. Embryos were then incubated for 3 h at 37° C.

### 2.3. Cryo-sections

After incubation, embryos were collected and fixed in 4% paraformaldehyde (PFA, in 1x PBS) at 4° C overnight. Excess tissue was then cut and excess PFA was removed by washing 3x with 1x PBS for 1 h. Embryos were cryoprotected by being incubated first in a 5% sucrose/PBS for 3 h and then in a 20 % sucrose/PBS solution at 4° C overnight, and were then embedded in a medium containing 7.5 % gelatin and 15 % sucrose dissolved in PBS, in a 35 mm Petri dish. Gelatin was left to set at room temperature and then stored at 4° C for up to a few days. Blocks were cut from these media and frozen using liquid nitrogen. 10 μm transversal slices were cut from these blocks using a cryostat (Reichert-Jung Cryocut E) at −21° C and placed onto gelatinized microscope slides (0.5 % gelatin, 0.05% chrome alum in water).

### 2.4. Immunofluorescence

Indirect immunofluorescence on the slices was performed following (Stern and Holland, 1993). All antibodies were diluted in a 1 % BSA/PBS solution. Primary antibodies used were rabbit anti-MARCKS (1:1000, Toledo et al., 2013), rabbit anti-cleaved Caspase 3 (1:200, Cell Signaling Technologies #9661), rabbit anti-ɣ tubulin (1:100, Sigma-Aldrich #T5192) and mouse anti-ZO-1 (1:50, Invitrogen #33-9100). Secondary antibodies used were anti-rabbit Alexa 488 (1:500, Invitrogen #A21206) and anti-mouse Alexa 488 (Invitrogen #A11001). Phalloidin-TRITC (1:8000-10000, Sigma) was included during the incubation with secondary antibodies. Hoechst 33342 DNA staining was performed during the final PBS washes.

### 2.5. Image Acquisition and processing

Slices were mounted for imaging using 70 % glycerol in Tris buffer (20 mM, pH 8), covered with a coverslip and sealed using nail polish. Imaging was performed on a confocal Zeiss LSM 800 microscope using a 20x (dry, NA = 0.8) or 25x (glycerol immersion, NA = 0.8) objective lens. Images consisted of stacks of optical sections every 1 μm, in slices corresponding approximately to the fourth somite. Image processing and analysis was done using FIJI (Schindelin et al., 2012).

### 2.6. Cell extrusion and cell death quantification

In order to determine the number of embryos exhibiting cell extrusion and indications of cell death (pyknotic nuclei and cleavage of caspase 3), mounted slices were examined under a Nikon Microphot FXA epifluorescence microscope, using 20x (dry, NA = 0.75) and 40x (dry, NA = 0.95) objective lenses. Extruded cells were identified as lying apical to the actin accumulation line corresponding to the apical side of the neuroepithelium in normal conditions, occasionally checking nuclear staining for confirmation. In order to determine that an embryo had not experienced any cell extrusion, four different nearby slices were checked. Counting of pyknotic nuclei and cells showing cleaved caspase 3 (i.e.: those labelled with the anti-cleaved caspase 3 antibody) was done in four nearby (non-damaged) slices of each embryo and adding the resulting numbers.

### 2.7. Protein distribution quantification

Starting with stacks of optical sections, distribution of fluorescent signals was determined from average intensity projections of three slices. Our region of interest was the mostly flat section of the neural plate adjacent to the medial hinge. Images were then rotated so as to make the apical side of our region of interest face left, and from there, the fluorescence profile was obtained along an approximately 20 μm wide transect (71 px, corresponding to 20.09 μm). Background signal intensity was obtained as the mean intensity of a 40 px square outside the tissue and subtracted from all values.

Phalloidin intensity profiles in control embryos typically showed two peaks corresponding to the basal and apical side of the tissue respectively, which we used to define these regions in the apico-basal axis. The apical region was defined as the interval within 9 px (2.5 μm) of the appropriate phalloidin signal maximum, which was observed to correspond approximately to a 50 % decrease in signal intensity in control embryos. The sub-apical region was defined as the interval between the basal phalloidin maximum and the apical region. In cases with cellular extrusion more apical peaks were observed, but a peak corresponding to the apical side of non-extruded cells was usually identifiable. Analysis of the tissue morphology was used to clear any doubts regarding multiple possible basal or apical peaks.

The ratio of apical to sub-apical signal for MARCKS and ZO-1 immunostaining was then calculated as the largest value found in the apical region divided by the average value in the sub-apical region.

### 2.8. Statistical analysis

Statistical analyses were performed for the values for the ratio of the maximum immunofluorescent signal intensity in the apical regions relative to the average signal intensity of the sub-apical region for both MARCKS and ZO-1 for each experimental condition, using GraphPad Prism (version 8.0.2 for Windows; GraphPad Software, Boston, Massachusetts USA, www.graphpad.com). All distributions passed the Shapiro-Wilk, Anderson-Darling and D’Agostino & Pearson lognormality tests (⍺ = 0.05), with the exception of the MARCKS distribution in control embryos, in which N was too small for the Anderson-Darling and D’Agostino & Pearson tests. Differences of means between pairs of experimental conditions were determined using Welch’s t-test for the natural logarithm of the presented values.

### 2.9. Theoretical description

We based our model on the augmented vertex model previously described by Kim et al. (2021, 2024), in which cell-cell interfaces are represented by vertices joined by straight edges, and nuclei are represented as solid-like objects constrained by the cell membrane. While this kind of model has usually been used to represent the apical surface of epithelia, our work represents the epithelium as a lateral cross-section, similar to those presented in the experimental results. Our 2D model is based on an energy functional (Eq. 1) depending on all cell areas, the length of all cell-cell (or cell-exterior milieu) interfaces and the total “overlapping” radial distance of a cell’s nucleus with respect to its membrane, integrated along the cell’s contour.

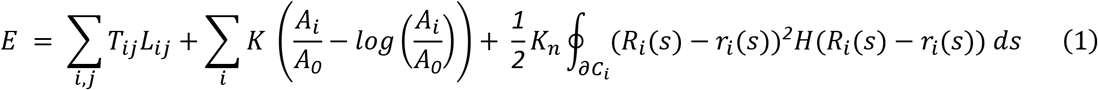

where *L*_*ij*_ and *T*_*ij*_are the length and the surface tension of the interface between cells *i* and *j* respectively, *K* is the cell compressibility modulus which measures how much pressure the cell exerts while being compressed, *A*_*i*_ is the area of cell *i, A*_*0*_ = *K*/*P*_*0*_ is a characteristic area based on *P*_*0*_ the external medium’s pressure, *K*_*n*_ is the nuclear elastic modulus which determines the repulsion force generated when the nucleus overlaps the cell boundary, *r*_*i*_(*s*) is the distance measured from the center of the nucleus of cell *i* to a point in the cell boundary *∂C*_*i*_ (parametrized by arc-length *s*), *R*_*i*_(*s*) is the distance between the nuclear center and its border in the same direction, and *H*(*x*) is the Heaviside function, which ensures only points in which the nucleus overlaps its cell’s membrane are included in the line integral (taken over cell’s *i* membrane *∂C*_*i*_). The first sum is over all cell-cell (and cell-exterior) interfaces *ij*, the second one over all cells *i*. Note that in the case of a circular nucleus, *R*_*i*_(*s*)is simply the nucleus’ radius (see a graphical description in Figure 1).

**Figure 1:**
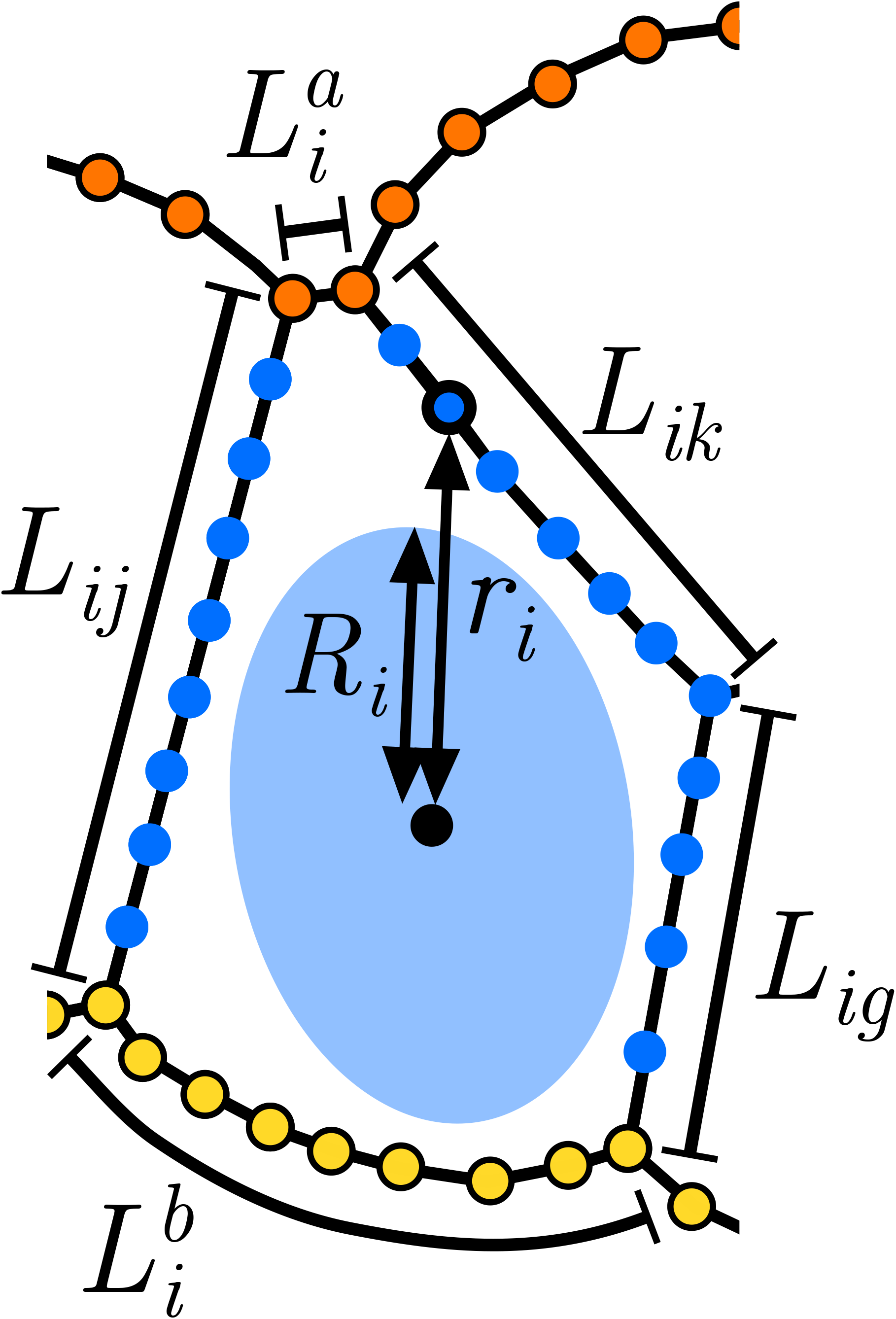
Diagram of terms included in the energy equation of our model. Here the vertices defining the boundary of cell i are represented along with its nucleus and its neighboring cells, similar to our simulation results. The first energy term in Eq. 1 depends on interface length *L*_*ij*_ which depends on the cells involved in the interface (e.g.: ij, ik, ig, as shown here). Apical and basal surfaces are here composed of yellow and orange vertices, with lengths marked as 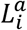 and 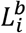 respectively. The third term in Eq. 1 corresponds to interactions with the nuclei and depends, for every point in the cell boundary, on its distance from the nuclear center *r*_*i*_ and on *R*_*i*_, the “radius” of the nucleus in the same direction. Note that this third term is only non-zero when *R*_*i*_ > *r*_*i*_, i.e.: when the nucleus overlaps the cell boundary.

### 2.10. Simulations

All simulations were based on a non-dimensionalized version of *E* (Equation 1; Appendix A.1) defining certain characteristic scales: *P*_*0*_ (pressure of the external medium), *T*_*0*_ (surface tension of cell-cell interfaces), 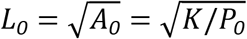 (characteristic length-scale) and *τ*_*R*_ (characteristic timescale for vertex relaxation). When not explicitly stated, we followed Kim et al. (2024) in their estimation for all corresponding parameters. Assuming a viscous medium where frictional forces dominate, the positions of all vertices follow first-order differential equations in which velocities are proportional to the sum of all non-viscous external forces (Odell et al., 1981). These forces may be obtained as the negative gradient of *E* with respect to their respective positions. The same can be applied to nuclei, which are defined as solids by their centers of mass and rotation angles, with this last variable being ignored by considering only circular nuclei in all simulations.

Dynamic simulations were carried out using custom scripts written for GNU Octave (Eaton et al., 2020) by numerically integrating these first-order differential equations using the Euler method with a time step *Δt* = *0*.*005 τ*_*R*_. Unless stated otherwise, all simulations were initialized with periodic boundary conditions of length *λ* = *10* and with *N* = *20* rectangular cells of area 1, a limit of our computational implementation. Edge length was maintained by adding or joining vertices between 1/16 and 1/8 of the average cell width, defined as *b* = *λ*/*N*. In the cases in which this resulted in the formation of a 4-fold vertex, a T1 transition (neighbour exchange; necessary for extrusion in our model) was evaluated and computed using the method described by (Duclut et al., 2021). The surface tension of all apical and basal interfaces was defined as *T*_*a*_, relative to *T*_*0*_. Whether the system had reached an equilibrium state was determined by qualitatively checking whether the energy had stabilized to a minimum over time.

In addition to these dynamical simulations, we analyzed the dimensionless energy minima for a simplified tissue made up of wedge-shaped cells without nuclei under different values of parameters *T*_*a*_/*T*_*0*_ and *b* (inversely related to the tissue’s cell density).

## 3. Results

### 3.1. Experimental assessment of the mode of apical cell extrusion in PMA-treated embryos

With the aim of elucidating if the PMA-induced apical cell extrusion in the neural plate was live or apoptotic extrusion, we set up the conditions for treating neurulating chick embryos simultaneously with PMA and the pan-caspase inhibitor QVD-OPh. Since both drugs must be administered solubilized in DMSO, and given the relatively low dilution rate needed for QVD-OPh, we used the lowest possible amount of DMSO which did not cause obvious effects on embryo or cell survival: 0.93%. In these conditions, control embryos developed normally, not showing delays in general development or neurulation (Fig. 2A). Widespread apical cell extrusion was observed throughout the neural plate in all embryos treated with PMA (n = 20) and those treated with both PMA and QVD-OPh (n = 20), while no cell extrusion was observed in control embryos (n = 15) or those under QVD-OPh treatment alone (n = 17) (Fig. 2B).

**Figure 2:**
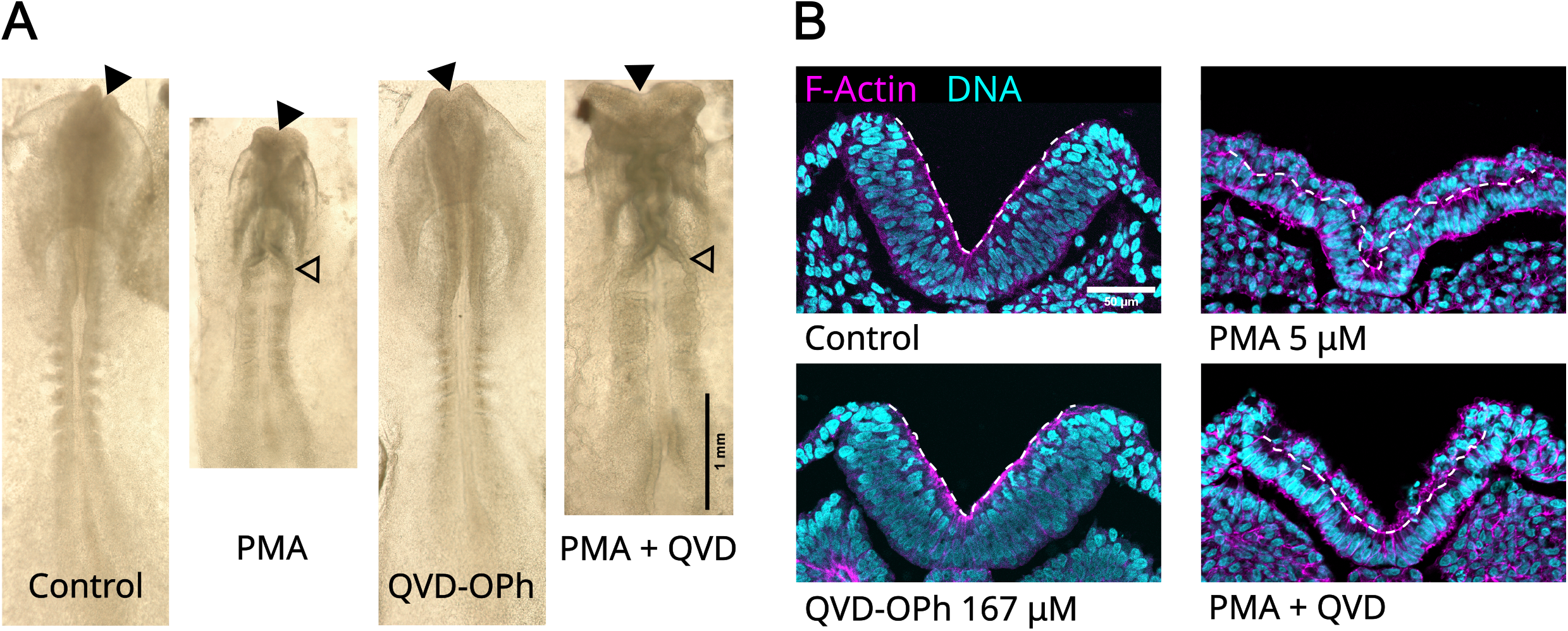
Comparison of chick embryos after treatment with PMA and QVD-OPh. **A**. Macroscopic view of treated embryos. Embryos treated with the pan-anti-caspase drug QVD-OPh seemed to develop similarly to Control embryos. Those treated with PMA, both alone and together with QVD-OPh, showed several defects, as described previously (Aparicio et al., 2018), including decreased length, neural tube closure failure and abnormally open anterior (filled arrowheads) and posterior (hollow arrowheads) neuropores. Scalebar: 1 mm. **B**. Transverse sections of the neural plate and adjacent tissues. Control and QVD-OPh-treated embryos displayed a thick pseudostratified epithelium. Treatment with PMA (including double treatment with QVD-OPh) produced widespread apical cell extrusion, evidenced by the presence of cells beyond the line of F-actin accumulation (dashed line; corresponding to the apical side of the layer of cells attached to the basal lamina), which are present along all regions of the neural plate. Scalebar: 50 μm

As expected, PKC activation with PMA caused in all cases a redistribution of MARCKS from the plasma membrane to the cytoplasm, in addition to a loss of its apical accumulation (Fig. 3A). QVD-OPh-only treated embryos, on the contrary, showed a MARCKS distribution that resembled control embryos. To further check the effect of these treatments on apico-basal polarity, we decided to analyze the distribution of the sub-apical cell adhesion–associated protein ZO-1. As previously reported (Aparicio et al., 2018), PMA treatment provoked a clear loss of apical accumulation of this apical marker, and this was not affected by co-incubation with QVD-OPh (Fig. 3B). In order to compare apical protein accumulation across the different treatments, we measured the ratio of apical signal with respect to the average sub-apical signal intensity for both proteins, MARCKS and ZO-1 (see section 2.7). QVD-OPh-only-treated embryos showed a similar distribution to control embryos, while those treated with PMA exhibited a significant decrease in apical protein accumulation (Fig. 3C). Similarly, double treated embryos showed a significant difference with respect to embryos treated only with QVD-OPh. These results further indicated that treatment with the anti-apoptotic drug did not affect cell extrusion or cell polarity by itself.

**Figure 3:**
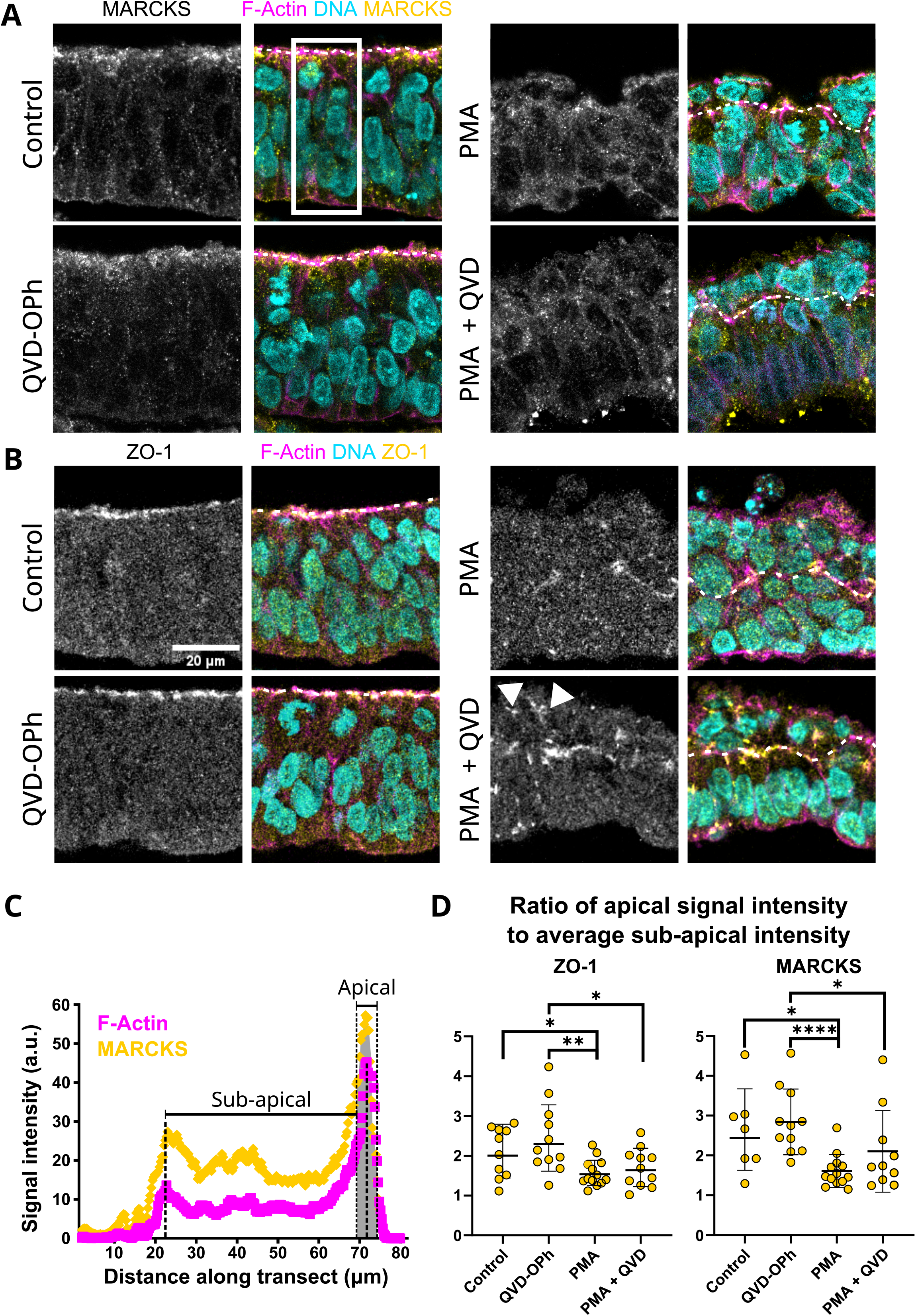
Distribution of MARCKS and ZO-1 along the apico-basal axis of the neuroepithelium. Intracellular distribution of MARCKS (**A**) and ZO-1 (**B**) in different experimental conditions. Both proteins accumulate near the apical side in control embryos, but have seemingly altered distributions under PMA and PMA + QVD-OPh double treatment, sometimes even presenting anomalous accumulations away from the line of F-actin accumulation (arrowheads). **C**. Example of intensity profiles used to determine the fluorescence intensity around the apical border (in gray; defined as the area of maximal F-actin accumulation) and along the sub-apical portion of the neuroepithelium in different conditions (only control condition example is shown). **D**. Plots for the ratio of the maximum immunofluorescent signal intensity in the apical regions relative to the average signal intensity of the sub-apical region for both proteins. Data shows significant differences between the geometric mean values for PMA-treated embryos with respect to Control (p-values = 0.0416 (ZO-1); 0.0279 (MARCKS)) and QVD-OPh-treated embryos (p-values = 0.0044 (ZO-1); <0.0001 (MARCKS)), and between QVD-OPh- and double-treatment embryos (p-values = 0.0233 (ZO-1); 0.0390 (MARCKS), indicating altered apico-basal polarity. Data are presented along with geometric mean and geometric standard deviation.

DNA staining indicated the presence of few scattered apparently dying cells in all embryonic tissues, at all treatment conditions. But, as previously described (Aparicio et al., 2018), PMA treatment greatly increased the number of pyknotic nuclei mostly on cells extruded from the neural plate (see Fig. 3B, for example). This was confirmed by immunolabeling of activated Caspase 3 (Fig. 4A). Most of these dying cells were observed beyond the apical border of the mass of extruded cells, indicating they were detaching from the epithelium. The double treatment of embryos with PMA and QVD-OPh, on the other hand, showed a sharp reduction in cell death as assessed by quantifying both parameters, demonstrating that the observed apical extrusion happens independently of cell death in these embryos, and can be defined as live cell extrusion (Fig. 4B).

**Figure 4:**
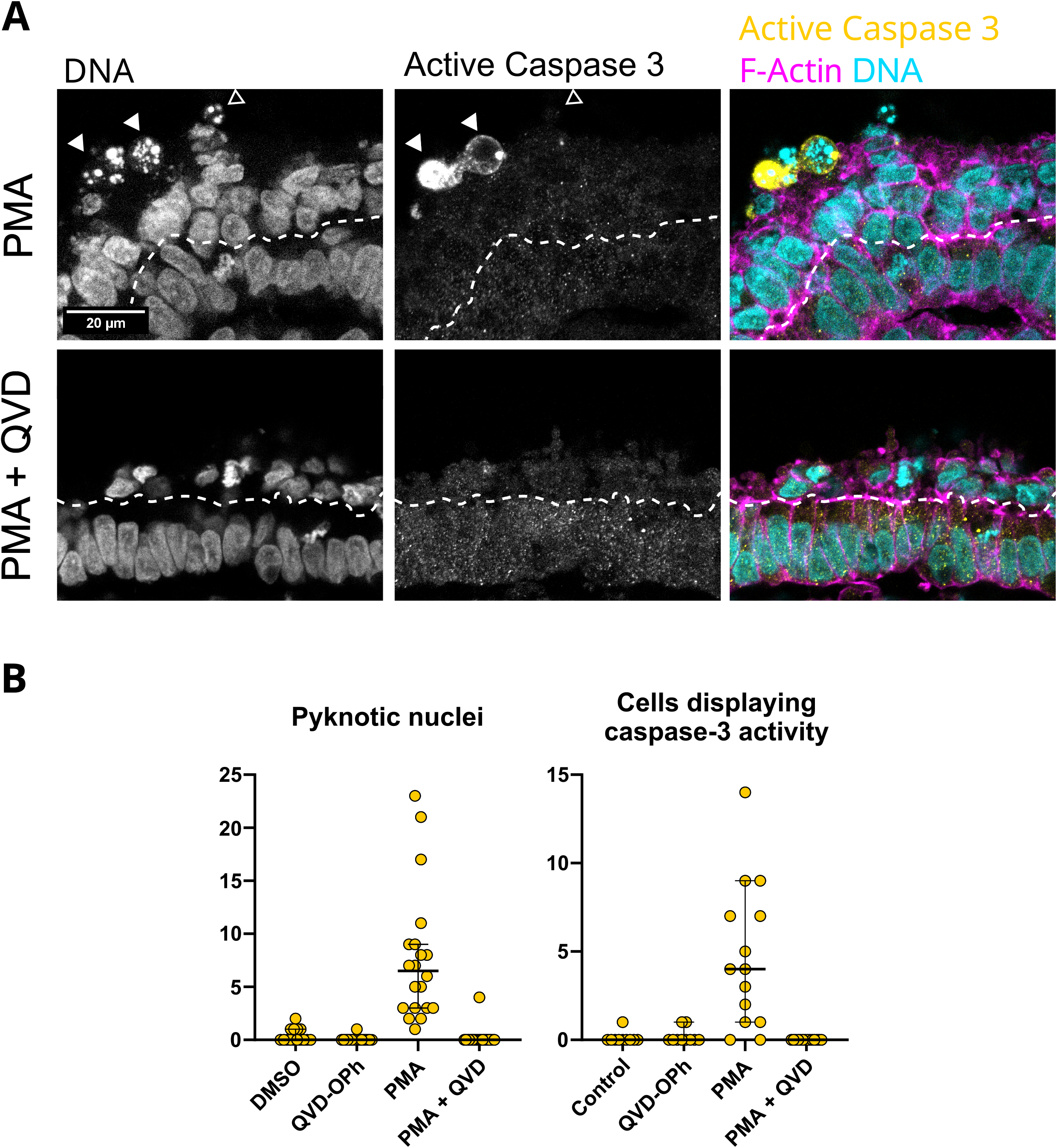
Occurrence of cell death among different experimental conditions. **A**. Examples taken from slices of embryos treated with PMA or PMA + QVD-OPh, illustrating the differences in signs of cell death. In PMA-treated embryos cells undergoing cell death were often observed, as indicated by the presence of pyknotic nuclei and/or caspase-3 activity (arrowheads). **B**. The number of pyknotic nuclei and cells displaying active caspase-3 immunodetection was higher in PMA-treated embryos than in all other experimental conditions. Most notably, double treatment with PMA and QVD-OPh abolished cell death while still inducing widespread cell extrusion. The dashed line marks the line of F-actin accumulation, corresponding to the apical side of the layer of cells attached to the basal lamina. Filled arrowheads: Cells exhibiting both pyknotic nuclei and active caspase-3. Hollow arrowhead: A cell with a pyknotic nucleus but no active caspase-3 immunoreactivity. Data presented along with median and 95% confidence interval.

### 3.2. Computational model of a pseudostratified epithelium and conditions leading to cell extrusion in the simulation

Since our experimental results support the idea that PMA treatment causes live cell extrusion during development, we focused on determining the conditions under which an epithelium like the neural plate could become mechanically unstable, leading to the elimination of live cells from the tissue. Our first goal was to produce a model which could reproduce the morphology of pseudostratified epithelia, that is, a single layer of columnar cells in which the nucleus occupies a large proportion of the cell volume, resulting in curved cell-cell interfaces shaped so as to accommodate all nuclei at different heights along the apico-basal axis. We are not considering in the model the continuous cell shape changes that would be caused by interkinetic nuclear migration in a living neuroepithelium because the experimental time-scale of the PMA-induced cell extrusion (less than 3 h) is much shorter than the average extension of the cell cycle in this cell population (8-12 h; Schoenwolf, 2018).

In order to achieve this, we developed a modification of the augmented vertex model originally described by Kim and collaborators to represent the dynamics of epithelia from an apical point of view (Kim et al., 2024, 2021). In our version of the model, we chose to represent a lateral view of an epithelial tissue under periodic boundary conditions (Fig. 5A). The model consists of a set of cells and solid-like nuclei for which an energy functional *E* is defined, which depends on the different cell areas, boundary lengths and position of the nuclei within the cell (see section 2.9 and Equation 1). Although we followed the force-based model of Kim et al. (2021, 2024) in defining this energy, the forces arising from our description are not necessarily identical to theirs, particularly in the case of the nuclear repulsion forces (for example, our calculations are based on vertices, not edges; see Appendix A.3). It can be proven that in the absence of nuclear repulsion forces, the angle between adjacent edges at equilibrium depends only on the interface they belong to (Appendix A.2). As a corollary, all interfaces not overlapping with a nucleus should approximate circular arcs at equilibrium. This indicates that an equilibrium state resembling a pseudostratified morphology could only be observed in cells with nuclei larger than the average cell width *b*, defined in this model as the dimensionless length of the tissue in the *x*-axis divided by the total number of cells. Supporting this, our initial dynamical simulations showed that cell layers containing no nuclei reached an equilibrium state consisting of identical columnar cells with straight cell-cell and convex cell-medium interfaces (Supplementary Fig. 1). This happened in less than twice the characteristic timescale for vertex relaxation *τ*_*R*_.

**Figure 5:**
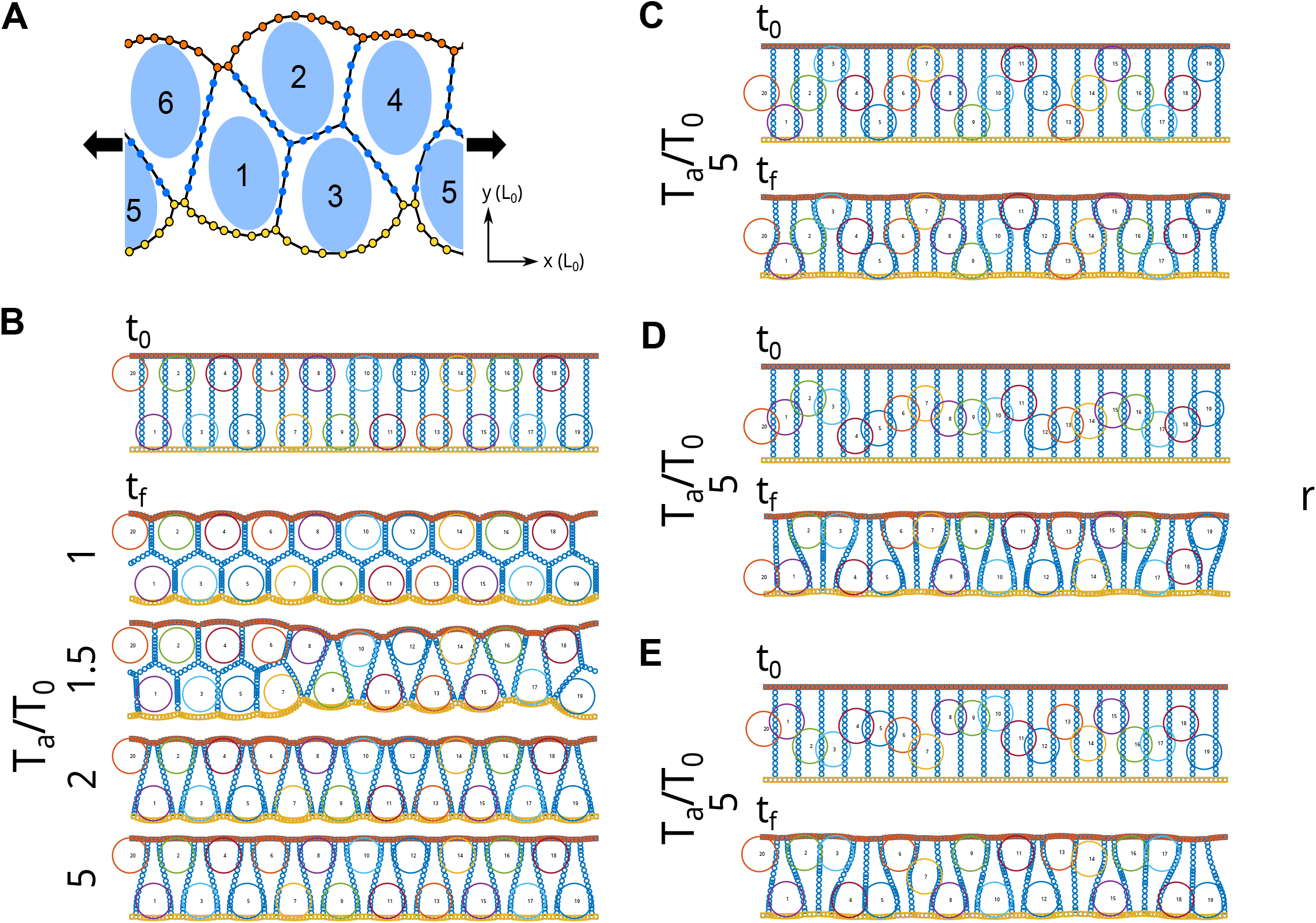
Dynamic simulations of pseudostratified epithelia including nuclei, using the energy model. **A**. Diagram depicting the state of a typical system at time t as obtained by dynamical simulation. Vertices are colored according to the type of interface they belong to (cell-cell interfaces are blue, while apical and basal interfaces are orange and yellow, respectively), and each cell contains a nucleus tagged according to the cell number. Periodic boundary conditions are used in the x-axis. Both axes are taken to be relative to the characteristic length scale 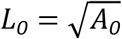. **B**. Long-term behavior of dynamic simulations in tissues with 20 cells, and an average cell width b = 0.5, under different values of *T*_*a*_/*T*_*0*_. All simulations start from a layer of identical rectangular cells with area *A*_0_ (= 1) and alternating nuclei, and are left to run for 20 characteristic time-lengths (*τ*_*R*_). These initial and final states are marked as t_0_ and t_f_, respectively. Energy profiles indicated an apparent equilibrium state for all cases except for when *T*_*a*_/*T*_*0*_ = *1*.*5*. For low values of *T*_*a*_/*T*_*0*_, a single layer morphology is unstable and cell extrusion ensues, while for larger values the single layer is maintained at equilibrium. For values larger than 5, a stable equilibrium solution resembling the pseudostratified epithelial morphology is reached. **C-E**. Simulations were performed at this stable value of *T*_*a*_/*T*_*0*_ = *5* for other nuclear configurations, all of which also proved stable in these conditions: a “staircase” distribution with intermediate nuclei (**C**) and different random nuclear distributions (**D**,**E**).

Based on the previous considerations, we followed by performing initial exploratory simulations including wide nuclei distributed alternatingly in the model, so that a cell with an “apical nucleus” (tangent to the initial apical boundary) has neighbors with “basal nuclei” and vice versa (Fig. 5B). Ratios of the apical/basal surface tension (*T*_*a*_) with respect to lateral surface tension (*T*_*0*_) of 1.5 or lower led to variable extensions of cell extrusion, with cells losing either their apical or basal interfaces, depending on their initial nuclear position, and stabilizing into a more rounded shape (Fig. 5B). Larger values of *T*_*a*_/*T*_*0*_, on the other hand, allowed for the tissue to stabilize into a single layer of columnar cells with curved cell-cell interfaces determined by the position of nuclei, corresponding to the pseudostratified morphology we sought (Fig. 5B). At values just above this threshold, cells formed a single layer but stabilized into a wedge-like shape with straight cell-cell interfaces, as would be expected in the absence of nuclei or with nuclei located in a region of the cell wider than itself (see Fig. 5B for *T*_*a*_/*T*_0_ = 2). Parameters *b* = *0*.*5, T*_*a*_/*T*_*0*_ = *5* prevented extrusion also in simulations initialized under different nuclear distributions (Fig. 5C-E).

### 3.3. Analysis of energetic minima in a simplified model

While the dynamical described simulations suggest a qualitative relation between relative surface tensions (*T*_*a*_/*T*_0_) and extrusion, we could not clearly express it in quantitative terms. Furthermore, we are limited by our computational implementation in the parameter ranges we can study, as well as by our lack of experimental measurements of these parameters in this particular tissue. For example, even though we know the neural plate is a highly dense tissue and suspect that this high density may be related to cell extrusion, we cannot perform dynamical simulations to directly test this for our system.

To avoid these limitations, we looked for general stability conditions by constructing a simplified version of our model for which we could better analyze equilibrium states and stability. Based on the results obtained from simulations with alternating nuclei, we considered a single-layered tissue formed by alternating wedge-shaped cells without nuclei (Fig. 6A; see Appendix A.4). We assume these cells to have continuous straight cell-cell boundaries (i.e.: no vertices), and their apical/basal interfaces to be circular arcs. Equations can be derived for such a tissue in terms of two geometrical variables: cell height (*h*) and basal width (*b*_*b*_), under parameters *T*_*a*_/*T*_0_ (apical/basal surface tension relative to lateral surface tension), *b* (average cell width, the inverse of cell density) and *L*_0_ (characteristic length, related to the cell compressibility module *K*). Setting *L*_0_ = 10 *T*_0_/*P*_0_ initially as in our dynamical simulations, we numerically computed the energy landscape for different cell shapes. Assuming the final area will be less than 1, we considered 0 ≤ *h* ≤ 1/*b* and 0 ≤ *b*_*b*_ ≤ 2*b*, between the limits where apical and basal extrusion occur. From these, we found the values of *h* and *b*_*b*_ which minimized the energy of the system for different values of parameters *b* and *T*_*a*_/*T*_0_, and searched for ways in which these might be related to extrusion or to single-cell layer stability.

**Figure 6:**
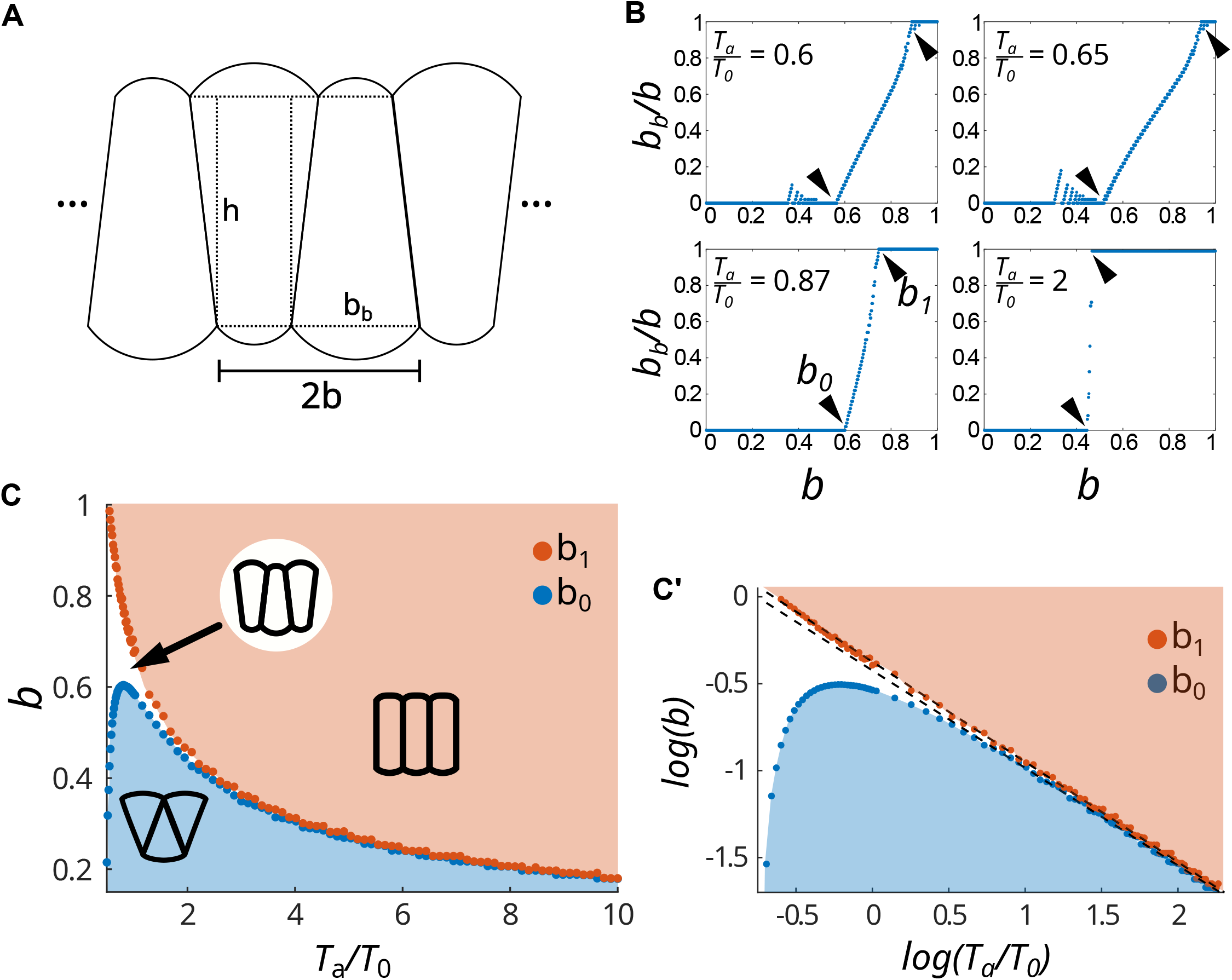
Long-term stability analysis for a monolayer composed of identical alternating wedge-shaped cells with no nuclei in terms of tissue parameters. **A**. To study the stability of a simple cell layer we consider an infinite tissue made up of alternating, identical wedge-shaped cells. The shape of one of these cells may be determined by its height (*h*) and its basal width (*b*_*b*_), assuming that the apical and basal interfaces correspond to circular arcs and that cell-cell interfaces are straight. The width of any two adjacent cells adds up to double the average cell width *b*, which is the inverse of cellular density. For each set of parameters, we find the basal width and height that minimize the total tissue energy. **B**. Value of the basal width *b*_*b*_ (expressed as a fraction of average cell width *b*) corresponding to the energy minimum vs *b*, here represented for different values of the ratio between apical/basal and lateral surface tension *T*_*a*_/*T*_*0*_. Although for low values of *T*_*a*_/*T*_*0*_ these graphs seem more complex, for values larger than 0.87, the plot of *b*_*b*_/*b* vs *b* seems to be divided into three distinct ranges: a low range where *b*_*b*_ = *0*, a middle range, and a high range where the energy minimum corresponds to *b*_*b*_ = *b*. Values *b*_*0*_ and *b*_*1*_ limit the low and high ranges for each *T*_*a*_/*T*_*0*_ (arrowheads). **C**. Plot of *b*_*0*_ (blue dotted line) and *b*_*1*_ (orange dotted line) as functions of *T*_*a*_/*T*_*0*_. For sufficiently large *T*_*a*_/*T*_*0*_, systems with parameters lying in the region above the *b*_*1*_ curve have their energy minima when *b*_*b*_ = *b* (epithelial stability) while those below *b*_*0*_ tend to *b*_*b*_ = *0* at equilibrium, which leads to cell extrusion. C’ shows the natural log-log plot of *b*_*0*_ and *b*_*1*_ vs. *T*_*a*_/*T*_*0*_. Fitted (dashed) lines correspond to equations *log*(*b*_*0*_) = −*0*.*5612 ln*(*T*_*a*_/*T*_*0*_) − *0*.*4237* and *log*(*b*_*1*_) = −*0*.*5761ln*(*T*_*a*_/*T*_*0*_) − *0*.*3729*. When simulations yielded values of *b*_*b*_/*b* larger than 1, they were replaced by *2* − *b*_*b*_/*b*, making use of the tissue apico-basal symmetry.

We observe that this system is symmetrical in that a tissue with basal width *b*_*b*_ is equivalent to one with basal width *2b* − *b*_*b*_, and that an energy minimum with *b*_*b*_ = *0* would lead either to extrusion or to a persistent 4-fold vertex, usually considered to be mechanically unstable. In general, low values of *b* lead to energy minima with *b*_*b*_ = *0*, and high values of *b* lead to *b*_*b*_ = *b*, with an intermediary transition range (Fig. 6B). For high values of *T*_*a*_/*T*_*0*_, this creates an apparently clear division of the *b* range into three connected regions, but this is not the case for lower values (*T*_*a*_ < *0*.*8 T*_*0*_, approximately), for which deviations may be observed. For example, low values of *b* may have a non-null equilibrium basal width *b*_*b*_.

The cases in which *b*_*b*_ = *b* correspond to a symmetrical equilibrium state which may be found in simulations when cells do not interact with the nuclei (Supplementary Fig. 1). This state always exists as a possible equilibrium solution, as equations for it can be found for all values of *b* as long as *T*_*a*_/*T*_*0*_ > *0*.*5* (Appendix A.5). We may define *b*_*0*_(*T*_*a*_/*T*_*0*_) as the maximum value of *b* for which the energy minimum occurs at *b*_*b*_ = *0* (or *2b*, equivalently) and *b*_1_(*T*_*a*_/*T*_*0*_) as the minimum value of *b* for which the energy minimum occurs at *b*_*b*_ = *b*.

Plotting both *b*_*0*_ and *b*_*1*_ against *T*_*a*_/*T*_*0*_ divides the parameter space into three regions according to the different long-time behaviours of the system (Fig. 6C). The upper region corresponds to *b*_*b*_ = *b* (stable monolayer composed of columnar cells of constant width), whiled the lower region corresponds to *b*_*b*_ = *0* (leading to extrusion). The distance between them tends to 0 for large *T*_*a*_/*T*_0_ values. We observe that *b*_*1*_(*T*_*a*_/*T*_*0*_) follows a power-law equation (Fig. 6D), and that the same can be said of *b*_*0*_(*T*_*a*_/*T*_*0*_) for sufficiently large values of *T*_*a*_/*T*_0_, obtaining equations

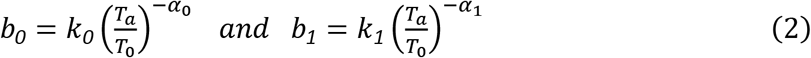

with *k*_*0*_ = *exp*(−*0*.*4237*) ≈ *0*.*65, α*_*0*_ = *0*.*5612* (linear fit of log-log data with *R*^*2*^ > *0*.*998* for *T*_*a*_/*T*_*0*_ > *1*.*6842*) and *k*_*1*_ = *exp*(−*0*.*3729*) ≈ *0*.*69, α*_*1*_ = *0*.*5761* (linear fit of log-log data with *R*^*2*^ > *0*.*999*). It should be noted that these values depend on the remaining parameter, the cell compressibility modulus *K*, with *k*_*0*_,*k*_*1*_ increasing and *α*_*0*_, *α*_*1*_ decreasing as *K* increases, seemingly reaching an asymptote (Supplementary Fig. 2). We could not find similar simple equations for other geometric parameters which might characterize the stability of the system, such as the cell height or aspect-ratio (*b*/*h*) at equilibrium.

### 3.4. Dynamic simulations of pseudostratified epithelia in conditions of near instability

Because of our simplification, this classification does not take into account the nuclear repulsion forces. Nevertheless, we may reasonably assume that for a system with nuclei to have a stable one-layered equilibrium state, it should also be stable in the absence of nuclei. Namely, in a highly dense tissue, where *b* is low and *h* is high, nuclei may be wider than *b* but much shorter than *h*, in which case they would only interact with a small number of vertices. If this approximation is correct, then the curve *b*_0_ should allow us to predict the threshold value of *T*_*a*_/*T*_0_ needed to stabilize a tissue for a given *b* even in the presence of nuclei. Our data show that *b*_*0*_ reaches a global maximum of *b* = *0*.*6040* between *T*_*a*_/*T*_*0*_ = *0*.*8* and *0*.*8167*, meaning: (1) that extrusion should not happen in any system with *b* > *0*.*6040* and (2) that every value *b* < *0*.*6040* has two possible threshold values for extrusion. We tested both situations by performing dynamical simulations using *b* = 0.5 and *b* = 10/16 = 0.625 (Fig. 7A).

**Figure 7:**
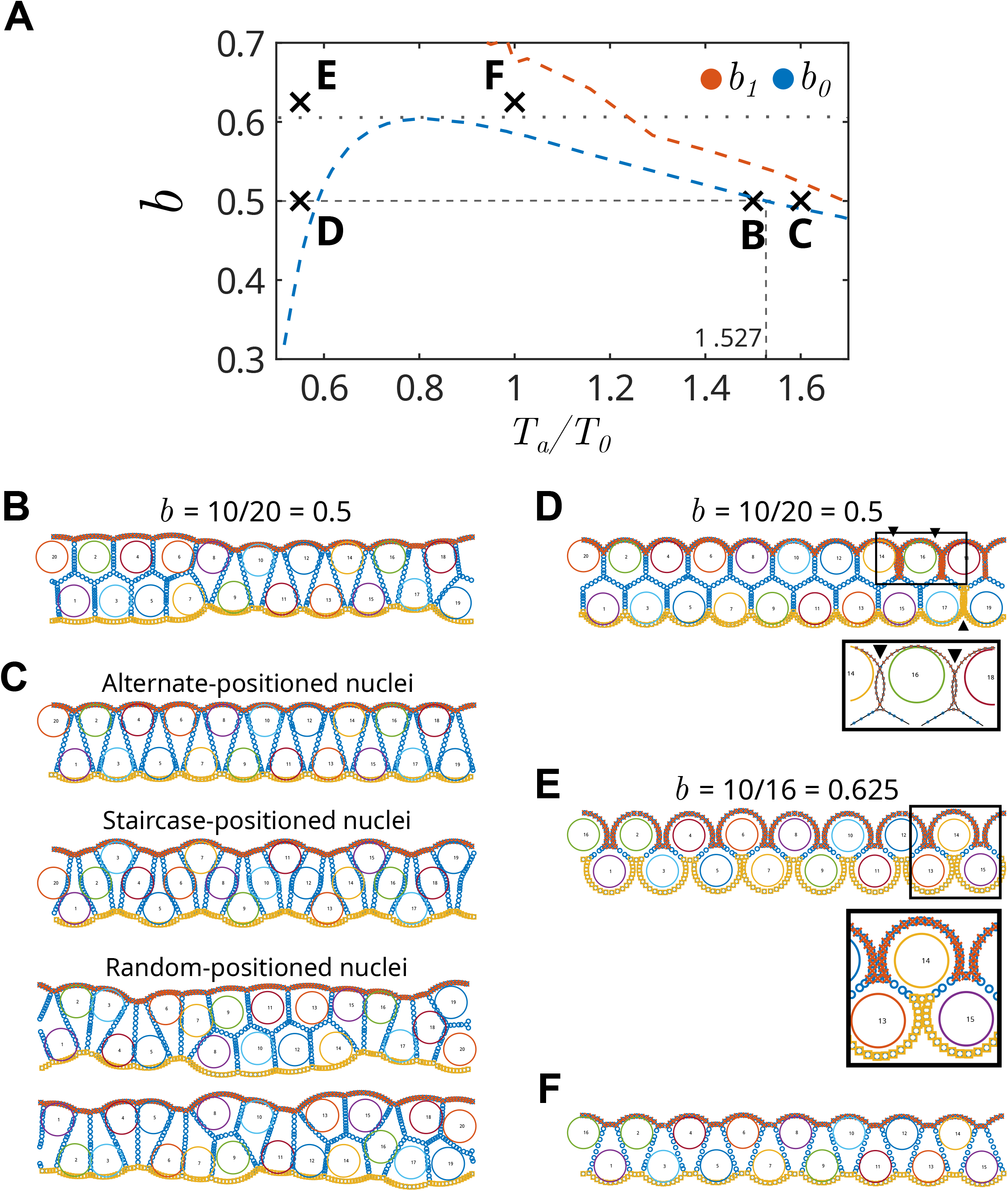
Comparison between dynamical simulations and stability predictions based on a tissue composed of wedge-shaped cells. Dynamical simulations like those of Fig. 5 were run here based on the quantitative analysis shown in Fig. 6, in and around the area in which wedge-shaped cells are expected in equilibrium, close to the stability threshold. **A**. Detail of *b*_*0*_ and *b*_*1*_ as functions of *T*_*a*_/*T*_*0*_, showing points corresponding to the simulations shown in B-F. Dashed gray lines point to the threshold value found for *T*_*a*_/*T*_*0*_ at *b* = *0*.*5*. The dotted gray line marks the calculated value of *b* = 0.604, above which there would be no cell extrusion. **B**. Starting with nuclei distributed alternatingly and *b* = *0*.*5*, we observe cell extrusion for *T*_*a*_/*T*_*0*_ = *1*.*5* (see Supplementary Video 1). **C**. Also at *b* = *0*.*5*, we can have many different outcomes for variable initial nuclear positions, at a tension ratio value that is slightly higher than the predicted threshold (*T*_*a*_/*T*_*0*_ = *1*.*6*). When nuclei start in an alternated position like in B or in a “staircase” distribution, cell extrusion is prevented (Supplementary Video 2). Random nuclear distributions, on the other hand, resulted in cell extrusion, as shown in the two bottom examples. **D**. In dynamical simulations with *b* = *0*.*5* and *T*_*a*_/*T*_*0*_ = *0*.*55*, not only did cell extrusion occur (contrary to our predictions) but also artefacts were introduced in the form of overlapping cell boundaries (arrowheads; detail in inset). **E**,**F**. Lowering cell density to 16 cells (*b* = *0*.*625*) means the system should lie above *b*_*0*_ according to our approximation regardless of surface tension. Although in these conditions and *T*_*a*_/*T*_*0*_ = *0*.*55* the cells adopt a rounded morphology, with a great expansion of one of their external surfaces, there is no cell extrusion (E). When the tension ratio was increased to *T*_*a*_/*T*_*0*_ = *1*, cells adopted a less-extreme wedge shape, also without extrusion as expected (F).

Our data predict that *b*_*0*_ = *0*.*5* when *T*_*a*_/*T*_0_ = 1.527 (Fig. 7A), meaning that systems with *b* = 0.5 should be unstable for *T*_*a*_/*T*_0_ ≤ 1.527 and stable for larger values, which agrees with our results for alternating nuclei (Fig. 7B,C; Supplementary Videos 1-2). Simulations using different initial nuclear distributions produced mixed results, with extrusion being prevented under a “staircase” distribution but not under random distributions (Fig. 7B) which had shown to be stable far away from this theoretical threshold, at *T*_*a*_/*T*_*0*_ = *5* (see Figure 5C-E). As mentioned above, there exists another possible threshold value for *b* = 0.5, meaning that extrusion should be prevented in a system with *T*_*a*_/*T*_0_ = 0.55. Nevertheless, extrusion was observed in this case along with simulation artefacts, indicating a limitation of our computational implementation (Fig. 7D). In agreement with our predictions, on the other hand, no extrusion was observed in the case where *b* = 0.625 when starting with an alternating distribution of nuclei, even for a low *T*_*a*_/*T*_*0*_ value (Fig. 7E,F). Taken together, these results partially validate our approximation but point to a dependence between the initial distribution of nuclei and the actual stability threshold.

## 4. Discussion

It was previously shown that MARCKS knockdown, or its phosphorylation upon PMA treatment, induced a loss of apico-basal polarity and extensive apical cell extrusion in the neural plate of chick embryos. This happened without an evident increase in cell death, except for cells that appeared completely detached from the epithelium, suggesting a live cell extrusion mechanism (Aparicio et al., 2018). In the present work, we have confirmed this to be the case: cell extrusion covering the full width of the neural plate occurred in all embryos under PMA treatment even when apoptosis was effectively inhibited with the anti-apoptotic drug QVD-OPh. Despite similar amounts of extruded cells, the treatment produced a near-total loss of signs of cell death. This allowed us to conclude that the observed apical cell extrusion occurred independently of apoptosis, indicating this to be an example of live cell extrusion that could be explained by mechanical alterations in the neuroepithelium (Eisenhoffer et al., 2012; Eisenhoffer and Rosenblatt, 2013).

Having cleared this point experimentally, we decided to proceed with a theoretical model of the neuroepithelium which could reproduce a pseudostratified morphology and show cell extrusion purely as a product of an imbalance of mechanical forces along the apico-basal cell axis. Previously published models, along with experimental studies, have pointed to losses in apical surface tension, tissue irregularities (i.e.: anisotropy and topological defects) and high in-plane cell densities as important conditions determining the occurrence of cell extrusion (Drozdowski and Schwarz, 2024; Marinari et al., 2012; Okuda et al., 2010; Saw et al., 2017). Excluding apical tension, the mechanical meaning of such predictors becomes difficult to discern in pseudostratified epithelia, where cell density is much higher than in previously considered situations and where cells develop complex three-dimensional shapes and arrangements in order to accommodate high nuclear densities (Gómez et al., 2021). Furthermore, these geometrical characteristics have been largely simplified or ignored by prior models of pseudostratified epithelia (Hecht et al., 2022; Ishii et al., 2021).

Although simulated extrusion may not perfectly reflect the *in vivo* mechanisms, we may use the model to find conditions under which a pseudostratified morphology might become unstable in a mechanical sense, leading to a live cell extrusion-like behavior. We found that a low ratio of apical/basal to lateral interfacial surface tension (*T*_*a*_/*T*_*0*_) led to widespread extrusion, while higher values prevented it. In particular, when wide nuclei (i.e.: wider than the average cell width) were present and *T*_*a*_/*T*_*0*_ was sufficiently large, the tissue reached an equilibrium state which was reminiscent of a pseudostratified epithelium: a single layer of cells with curved boundaries shaped so as to accommodate all nuclei at different heights along the apico-basal axis.

With these considerations in mind, the results obtained from our model would suggest that, under normal conditions, the surface tension in apical and basal interfaces must be larger than that in lateral cell-cell interfaces so as to mechanically stabilize the pseudostratified morphology. To quantitatively assess this, we approximated the stability of cells in a dense tissue where the nucleus mechanically interacts with a small fraction of the cell’s surface by analyzing a simple tissue composed of identical wedge-shaped cells without nuclei. Under this simplification, we found that the minimum average cell width *b*_*0*_ required for a tissue with relative apical/basal surface tension *T*_*a*_/*T*_*0*_ to be stable approximately follows a power law, with parameters depending on the cell compressibility modulus *K*. This approximation, while correct for large values of *T*_*a*_/*T*_*0*_ or *b* and certain regular nuclear distributions, was not enough to predict cell extrusion in general, particularly when nuclei had random initial positions which had been shown to be stable under higher *T*_*a*_/*T*_*0*_ values. These results serve to point out the limitations of our approximation and show that the exact conditions for extrusion may depend on the actual nuclear distribution adopted by the tissue. Unlike alternating nuclei, some configurations require nuclear rearrangements to reach stability, particularly those with initially overlapping nuclei such as the random distributions tested here. In our model these rearrangements would alter the force balance on the cell boundary in complex ways, which may explain these differences in stability thresholds. The importance of nuclear positioning, as well as the forces it exerts on the cell cortex through interactions with the cytoskeleton, have been experimentally demonstrated in different epithelial morphogenetic processes (Ambrosini et al., 2019; Ferme et al., 2025; Roby and Rauzi, 2025).

Although the relative simplicity of this theoretical model does not reflect most of the complex features of a real epithelium, the concomitant inclusion of a surface tension and a non-deformable nucleus can be used to simulate morphogenetic processes in which both factors are important (Kim et al., 2024). Deformability is nevertheless an intrinsic property of living cell nuclei, although with some limitations (Kalukula et al., 2022). Interestingly, in the extension of the Drosophila embryo germband, the simultaneous genetic alteration of nuclear deformability and the pharmacological destabilization of microtubules, affecting nuclear positioning in the apico-basal cell axis, causes both morphogenetic defects and cell extrusion (de Leeuw et al., 2024).

Based on the previous analysis, we can give an interpretation of our experimental results simply by assuming that PMA treatment alters the relative surface tension of the apical and/or basal interface (e.g.: by diminishing apico-basal polarization in each cell) without altering cell compressibility. The change required to destabilize the single-layered morphology of the tissue would imply reducing *T*_*a*_/*T*_*0*_ so that the new minimum stable cell width *b*_*0*_^*PMA*^ became larger than the average cell width *b* of the tissue. Furthermore, although the previous approximation of the tissue as formed by wedge-shaped cells no longer applies to the whole tissue once extrusion begins, we may use it to approximate only the basal-most layer of cells, for which the main effect of extrusion is to reduce the cellular density, that is, to increase *b*. If extrusion eventually stops and the tissue reaches a new stable state, then the average cell in the basalmost layer should reach an expected final width greater than *b*_*0*_^*PMA*^. In this simplified explanation, both the occurrence and the extent to which cell extrusion happens should be mainly determined by the relative decrease in the value of *T*_*a*_/*T*_*0*_ due to PMA treatment, a consequence of the approximate power-law structure of the stability threshold.

When extrapolating these observations to the living tissue, we could assimilate polarized changes in surface tension to alterations in cell polarization (Moazzeni et al., 2025; Narayanan et al., 2023). Treatment with PMA causes a loss of apico-basal cell polarity, evidenced by a redistribution of the apical marker ZO-1, as well as of actin filaments. As expected, there was also a re-localization of the main PKC substrate MARCKS, previously shown to be in great part responsible for PMA-induced cell polarity downregulation and cell extrusion in the chick neural plate (Aparicio et al., 2018). Evidence from this previous work also indicated that the PMA effect on these cells could involve a destabilization of cortical actin filaments, probably leading to a general reduction in cell surface tension (Chugh et al., 2017). This cortical actin destabilization could be explained by an alteration in MARCKS function owed to its phosphorylation at the effector domain by PKC, blocking its ability of sequestering PIP_2_ at the plasma membrane (Laux et al., 2000). Different lines of evidence have linked MARCKS to the modulation of epithelial cell mechanical stability, and its involvement in processes such as morphogenesis and carcinogenesis (Veloz et al., 2021). All the mentioned experimental evidence points to an effect of PMA mostly on apical membrane tension, given the high apical accumulation of MARCKS and actin filaments in neural plate cells, and that the basal membrane domain is tightly associated to a rigid external scaffold, the basal lamina.

In summary, we have presented here evidence of the occurrence of live cell extrusion in the neuroepithelium during neurulation, which could be caused by a reduction in cortical actin cytoskeleton integrity and a concomitant down-regulation of apico-basal cell polarity. In order to find a way of explaining these results within a mechanical framework, we have introduced an expanded energy-based vertex model allowing for the presence of nuclei and curved interfaces which is able to represent pseudostratified epithelial morphology. In line with previous research, our work points to a high relative apical/basal surface tension as being the main stabilizing factor against cell extrusion in high-density tissues such as the developing neural plate. Furthermore, we derive a novel approximate condition for live cell extrusion relating cell density and surface tension, which follows a power-law equation. Measurements of surface tensions and cell densities in the developing neural plate could help validate these results and contrast our approximation with other similar pre-existing models.

## Acknowledgements

The authors gratefully acknowledge Dr. Lilián Perdomo and Prodhin SA for providing the fertilized hen eggs, and the Advanced Bioimaging Unit at the Institut Pasteur Montevideo for their support and assistance with microscopy. We thank Gonzalo Aparicio for initial help with chick embryo manipulation and fruitful discussion on the experimental aspects.

## Author contributions

SABR: Conceptualization, Data curation, Formal analysis, Investigation, Methodology, Software, Writing – original draft, Writing – review and editing; JAH: Conceptualization, Formal analysis, Methodology, Supervision, Writing – review and editing; FRZ: Conceptualization, Data curation, Formal analysis, Funding acquisition, Methodology, Project administration, Resources, Supervision, Writing – original draft, Writing – review and editing.

## Data availability statement

Original data and custom scripts written for GNU Octave are available at: Bosch, Santiago; Hernandez, Julio; Zolessi, Flavio (2025), “Data for “Mechanical conditions preventing live cell extrusion during primary neurulation in amniotes”“, Mendeley Data, V1, doi: 10.17632/pc7ytf2xt7.1. Other data and materials are available from the corresponding author upon reasonable request.

---

**Supplementary Figure 1:**
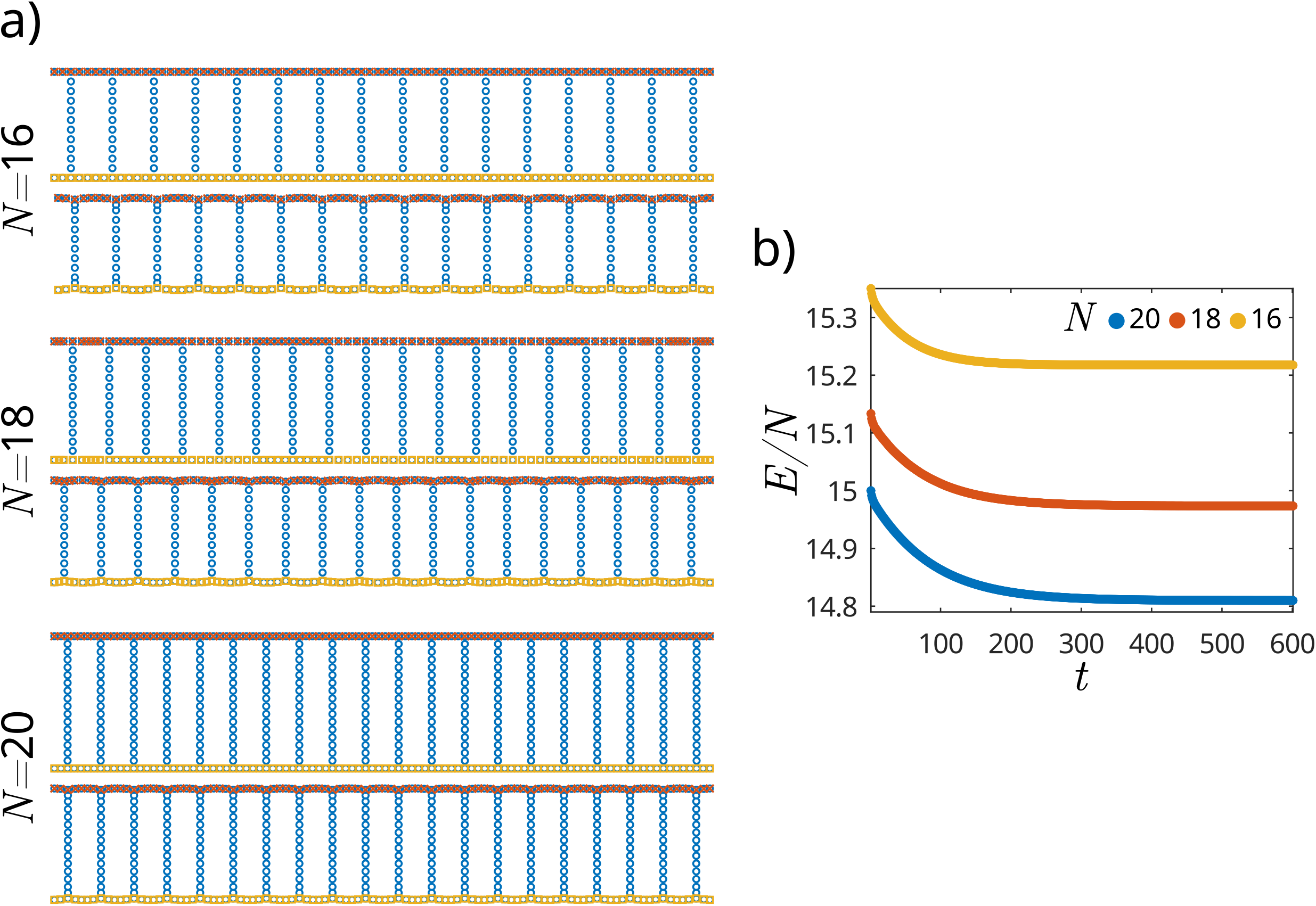
Initial simulations performed without nuclei quickly reach equilibrium in less than *21:*_*R*_. (A) Starting from an initial configuration of *N* identical rectangular cells with unit area, a stable state was reached composed of cells with straight cell-cell and convex cell-exterior interfaces, here pictured for *N* **=** *16, 18, 20* after 400 integration steps (equivalent to twice the characteristic time for vertex relaxation). This state was achieved even when the initial vertex distribution was not homogeneous, as in the case of N **=** *18*. (B) Plot of energy E per cell in time. Initial and final values were quantitatively similar for all cases, although the relaxation time became larger the smaller the average cell width b, with *t*_*50*_ (the time required to lose half the total lost energy) being 0.165*τ*_*R*_, 0.235*τ*_*R*_ and *0*.*27 τ*_*R*_ for *N* **=** *16, 18* and *20* respectively. Here we continued the simulation until 3rR, but we can qualitatively see that the system has reached equilibrium before that. *T*_*a*_*/T*_*0*_ = 3 for all shown simulations.

**Supplementary Figure 2:**
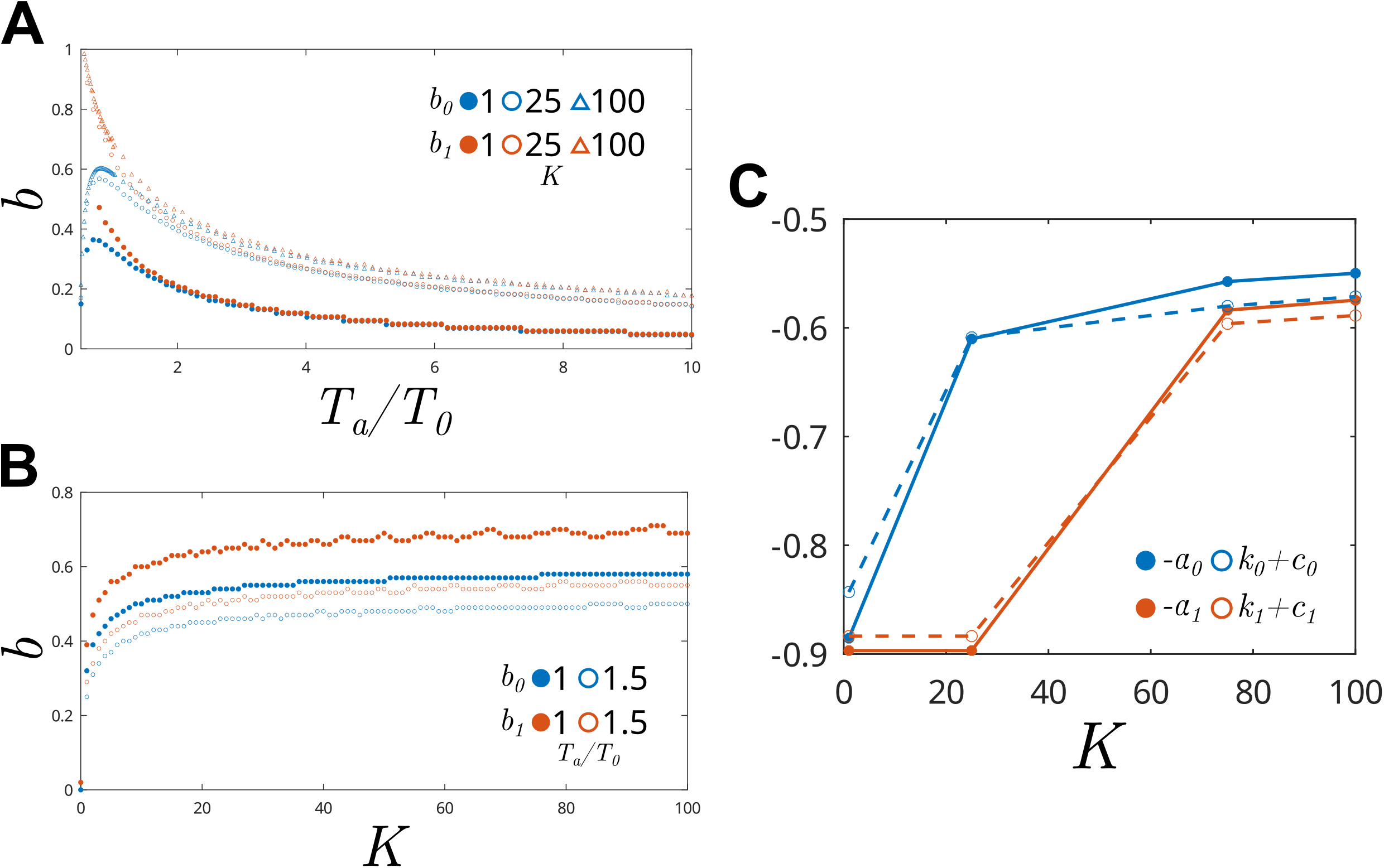
Dependence of curves *b*_*0*_ and *b*_*1*_ with parameter *K*. (a) Plot of b_*0*_(*T*_*a*_/*T*_*0*_) and *b*_*1*_ (*T*_*a*_/*T*_*0*_) for *K* = *1, 25, 100*. Increasing *K* makes *b*_*0*_ and *b*_*1*_ generally have larger values, although the difference becomes smaller for larger *K*. (b) Plot of *b0*(*K*) and *b1* (*K*) for *Ta*/*T0* = *1, 1*.*5*. As shown in (a), at each *T*_*a*_/*T*_*0*_, both *b*_*0*_ and *b*_*1*_ grow monotonically with *K*, seemingly reaching an asymptote at high *K*. (c) Plot of parameters *k*_*0*_, *k*_*1*_ (open circles, broken lines), *-a*_*0*_ and *-a*_*1*_ (closed circles, lines) as functions of *K*, obtained by fitting *b*_*0*_(*T*_*a*_/*T*_*0*_) and b_*1*_ (*T*_*a*_/T_*0*_) to a power-law equation *k*(*T*_*a*_/*T*_*0*_f^-*a*^. Because values for *a*_*0*_ and *k*_*0*_ seemed to be parallel to each other, in order to better visualize them together we defined *c*_*0*_ as the average difference between them, finding *c*_*0*_ = *-1*.*2125*. Correspondingly for *a*_*1*_ and *k*_*1*_, we found *c*_*1*_ = *-1*.*2756*.

**Supplementary Video 1:** Evolution in time of a simulated tissue in which extrusion occurs, as shown in Figure 7B. In this simulation the average cell width is *b* = *0*.*5* and *T*_*a*_/*T*_*0*_ = *1*.*5*, which leads to extrusion. Although initially the tissue morphology seems to stabilize, several T1 transitions happen at *t* = *11*.*8 τ*_*R*_, causing the state of the tissue to change greatly. At *t* = *20 τ*_*R*_ the system has not yet reached a new equilibrium. That the stability threshold for tissues with *b* = *0*.*5* is between *T*_*a*_/*T*_*0*_ = *1*.*5* and *1*.*6* is supported by the stability analysis performed on a simplified model.

**Supplementary Video 2:** Evolution in time of a simulated tissue in which extrusion does not occur, as shown in Figure 7C. In this simulation the average cell width is *b* = *0*.*5* and *T*_*a*_/*T*_*0*_ = *1*.*6*. The time it takes for the energy to relax 90% into its final value is less than *t* = *τ*_*R*_, but in this example the tissue qualitatively can be seen to stabilize in around *6 τ*_*R*_. That the stability threshold for tissues with *b* = *0*.*5* is between *T*_*a*_/*T*_*0*_ = *1*.*5* and *1*.*6* is supported by the stability analysis performed on a simplified model.

## Appendix A

### A.1 Non-dimensional version of energy functional

For all simulations and for the following derivations, a non-dimensional version of the energy functional *E* (eq. 1) was used. The formula can be obtained as *ϵ* = *E*/*T*_*0*_*L*_*0*_, with *T*_*0*_ being a characteristic surface tension equal to the surface tension of any cell-cell interface, and 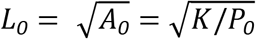 a characteristic length, where *P*_*0*_ is the pressure of the external medium. Using this, we obtain

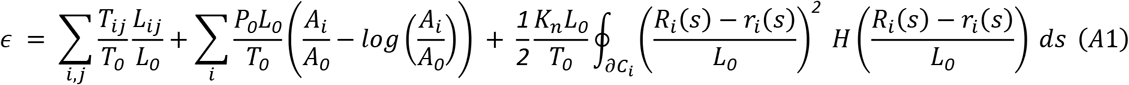

where all variables are relative to *L*_*0*_ and *A*_*0*_ and three dimensionless parameter groups (*T*_*ij*_/*T*_*0*_, *P*_*0*_*L*_*0*_/*T*_*0*_ and *K*_*n*_*L*_*0*_/*T*_*0*_) have been defined. We will use the dimensional and dimensionless names for these variables interchangeably throughout this appendix (i.e.: *E* for *ϵ, L*_*ij*_ for *L*_*ij*_/*L*_*0*_, etc.).

### A.2 Force balance and shape of an interface at equilibrium (without nuclear overlap)

Two types of forces act on all vertices which don’t interact with the nucleus: a tangential force coming from the surface tension term and a force derived from the pressure term in *E* (eq. 1). Let 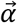 be a 2-fold vertex on the boundary between cells *i* and *j* (any of which we might take as the external medium, substituting *A* = *1*/*P*_*0*_), with clockwise neighbour vertex (*x*_−_, *y*_−_) and counter-clockwise neighbour (*x*_+_, *y*_+_). Then the force acting on 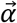 is given by:

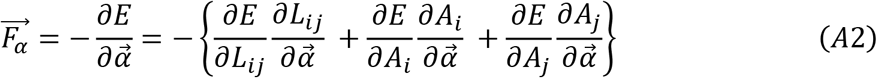

Because 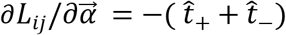 (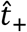 and 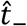 being the tangent vectors pointing away from vertex 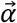) and 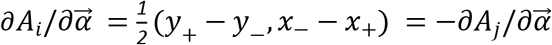 (Fletcher *et al*., 2013), we can rewrite equation (A2) as

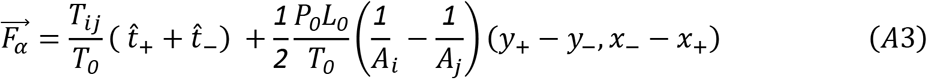

We observe that 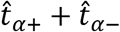 always points in the average direction between the two edges and that (*y*_+_ − *y*_−_, *x*_−_ − *x*_+_) is perpendicular to the line between (*x*_−_, *y*_−_) and (*x*_+_, *y*_+_). In equilibrium *F*_*α*_ = *0* and therefore the two terms in equation (A3) must be in the same direction (since *T*_*ij*_ ≠ *0*), meaning 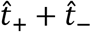 is perpendicular to (*x*_+_ − *x*_−_, *y*_+_ − *y*_−_). Since 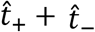 bisects the angle between both edges, it follows that both edges must have the same length *l*, and since (*y*_+_ − *y*_−_, *x*_−_ − *x*_+_) has the same length as (*x*_+_ − *x*_−_, *y*_+_ − *y*_−_), we may write

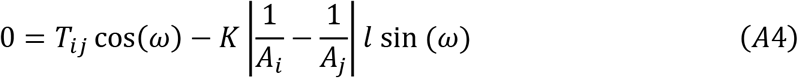

where *2ω* the inner angle between the two edges.

Since *T*_*ij*_ and |*1*/*A*_*i*_ − *1*/*A*_*j*_ | depend only on the cells involved, the inner angle *2ω* is only a function of edge length, which is equal between any two adjacent 2-fold edges and therefore constant along an interface. This means that any interface between two cells (or between a cell and the external medium) with no overlapping nucleus approximates a circular arc at equilibrium. Additionally as expected, the edge curves towards the cell with the least pressure (largest area).

### A.3 Deduction of nuclear repulsion forces and relation to the model by Kim *et al* (2024)

In vertices experiencing nuclear repulsion, 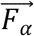 includes terms of the form

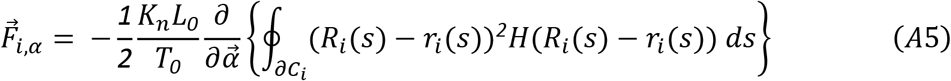

This derivative involves changes in both the path for the integral and the value of *r*_*i*_(*s*) along that path. The integral is only modified along subsequent straight paths *γ*_−_ and *γ*_+_, which go from 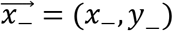 to 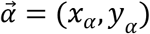 and from 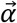 to 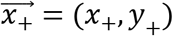, respectively. For example, the integral *I*_−_along *γ*_−_ can be written as

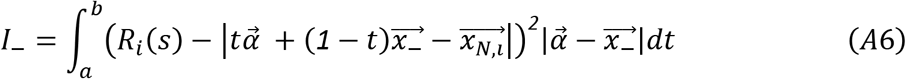

Where 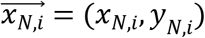 is the nuclear center, 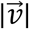 is the Euclidean norm, and where we have omitted the Heaviside function by only integrating in the interval where it is non-zero (*a* and *b* are both functions of 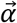 with values in the interval [*0,1*]). This integral is of the form 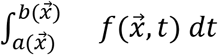 and therefore its derivative with respect to 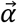 (i.e.: taken with respect to 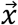 and evaluated at 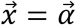) follows the Leibniz Integral Rule:

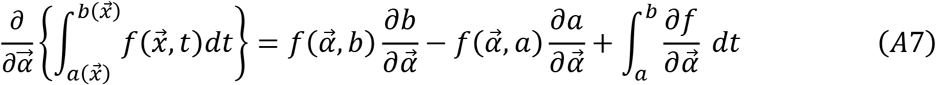

With 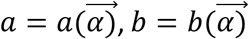. Since *a* is the lower limit where the (smooth) integrand of equation A6 is positive, either 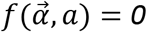 or *a* is the beginning of the edge (i.e.: *a* = *0*) in which case 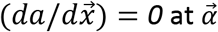. Applying a similar reasoning to *b* simplifies equation (A7) to

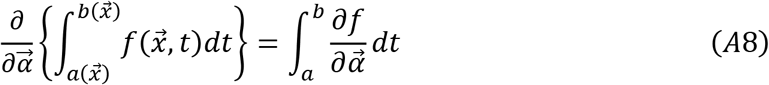

For simplicity, we assume circular/spherical nuclei –as in our simulations– and take *R*_*i*_(*s*) = *R*. Rewriting 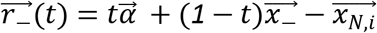, differentiation of the integral along *γ*_−_ (eq. A6) with respect to the position of vertex 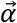 yields

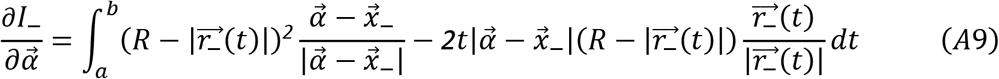

Defining 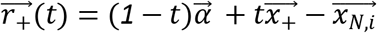 and redefining *a* and *b* appropriately, one may differentiate along *γ*_+_, resulting in

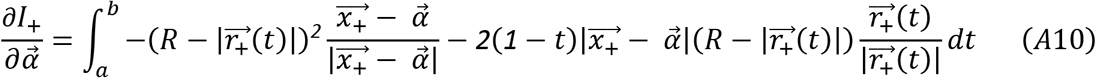

Finally, the nuclear repulsion force acting on vertex 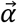 is proportional to the sum of both these integrals, that is

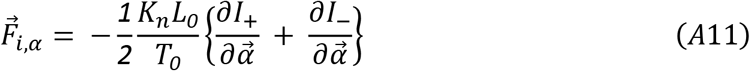

Our integral term in the energy functional *E* may be seen as a potential for the repulsive interaction included by Kim *et al*. (2024) in their force-based model. Although the force derived from our model is not equal to that proposed by them, we may state a similarity in the formula for the total force arising from the repulsion of a single edge. This is equivalent to the sum of the derivatives of a single edge’s path integral, differentiated with respect to the position of both of its endpoints. Using the integral along *γ*_−_ as an example, we obtain a force 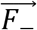 of the form

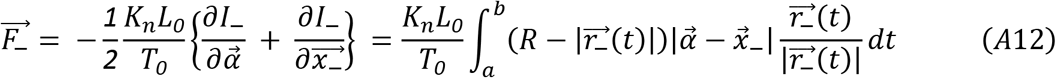

Re-introducing the Heaviside function, parametrizing by arc-length and writing the edge length as *L*_−_ we can write this as

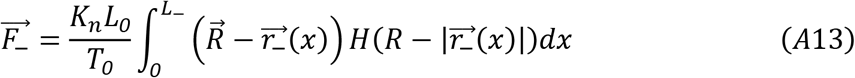

which follows the form proposed by Kim *et al*. 2024 as *F*_*αβ*_.

### A.4 Derivation of energy formula for wedge-shaped cells

Without nuclei, the dimensionless energy functional (eq. A1) becomes

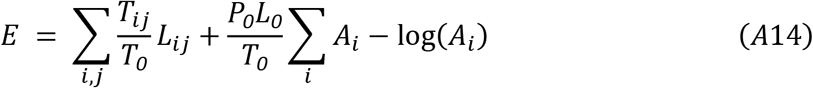

Therefore, all we need to determine its minima are formulas for the cell areas and interfacial lengths. Surface tension is *T*_*0*_ at cell-cell and *T*_*a*_ at apical/basal interfaces. The energy per cell is thus

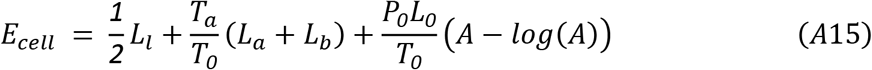

with *L*_*l*_, *L*_*a*_ and *L*_*b*_ the lateral, apical and basal interface lengths, defined as in Figure A1a. The average cell width is *b* = (*b*_*a*_ + *b*_*b*_)/*2*, equal to the sum of the apical (basal) widths of any two adjacent cells.

The apical/basal sides are circular arcs (Fig. A1b; see above) and we see that for a generic circular arc of base *b* and angle *θ* with respect to baseline, the radius of curvature is given by *ρ* = *b*/*2sin*(*θ*). The arc length is then given by

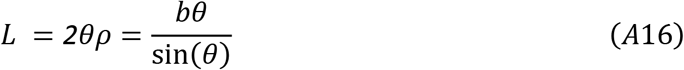

while the cap area is equal to

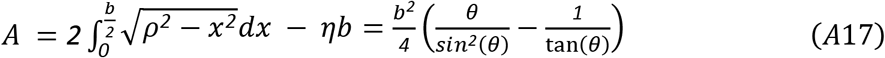

where the integral corresponds to the shaded area in Fig. A1b, and we have substituted *ρ* = *b*/*2sin*(*θ*), *ρ*^*2*^ = *η*^*2*^ + (*b*/*2*)^*2*^ and *η* = *b*/*2tan*(*θ*).

Finally, the interface lengths and total area are given by

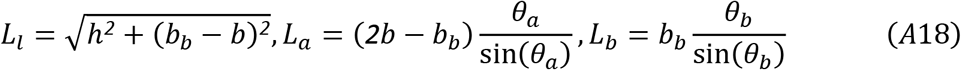

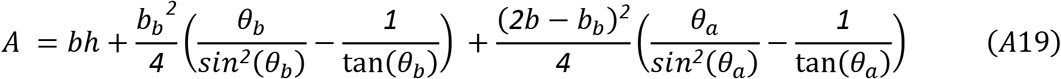

where we have substituted *b*_*a*_ = *2b* − *b*_*b*_ and *b*_*b*_ − *b*_*a*_ = *2*(*b*_*b*_ − *b*).

Although four variables (plus parameter *b*) appear in equations (A18) and (A19) (and consequently in *E*_*cell*_), *θ*_*a*_ and *θ*_*b*_ are actually functions of *h* and *b*_*b*_. This can be proven by considering a 3-fold vertex in the tissue, defined by angles *θ*_*a*_, *θ*_*b*_ and *𝜙* (Fig. A1c). In the limit in which inter-vertex distance tends to 0, pressure-derived forces disappear (see eq. A3, A4), so the tensions must add up to 0 in equilibrium, that is

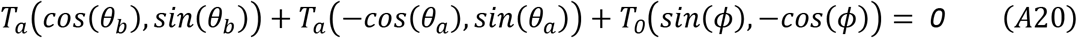

or equivalently

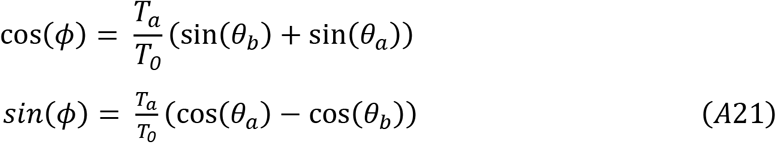

These equations can be solved by introducing variables *α* = (*θ*_*a*_ + *θ*_*b*_)/*2* and *β* = (*θ*_*a*_ − *θ*_*b*_)/*2*, so that

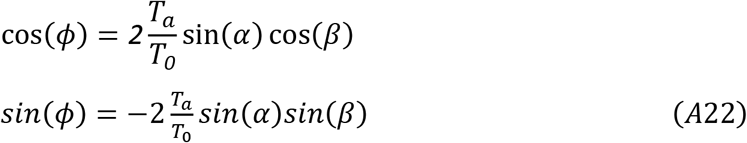

Dividing the second equation of (A22) by the first, we obtain

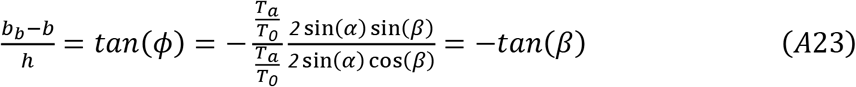

where we used the fact that *tan*(*𝜙*) = (*b*_*b*_ − *b*_*a*_)/*2h* = (*b*_*b*_ − *b*)/*h* (Fig. A1a). Because *θ*_*a*_, *θ*_*b*_ ∈ (*0, π*/*2*) and *θ*_*a*_ < *θ*_*b*_, there is only one value of *β* ∈ (−*π*/*2,0*) which satisfies equation A22. This means that *𝜙* = −*β*, which in turn simplifies the first equation in A21 to

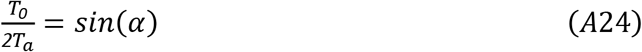

with angle *α* ∈ (*0, π*/*2*) being uniquely determined. From eq. A23 and A24 we obtain

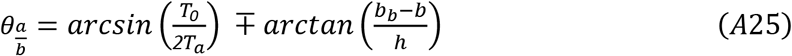

which proves that *E*_*cell*_ is a function of variables *h* and *b*_*b*_ only.

**Figure A1:**
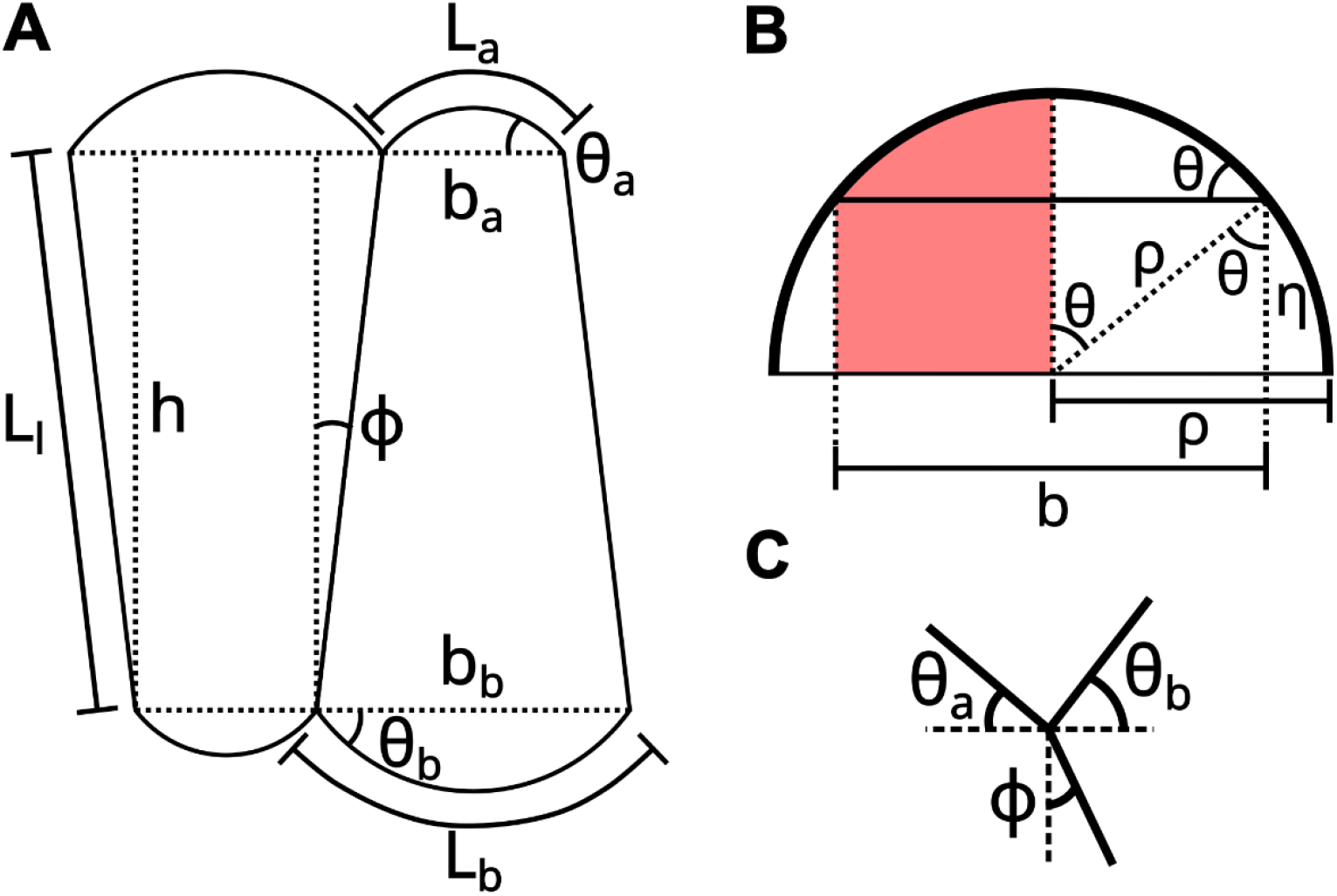
Diagrams for the derivation of cell area and interfacial lengths in the simplified tissue composed of wedge-shaped cells. **A**. Diagram of two neighbouring cells, indicating basal and apical widths (*b*_*b*_ and *b*_*a*_), contact angles (*θ*_*b*_ and *θ*_*a*_) and surface lengths (*L*_*b*_ and *L*_*a*_), as well as the lengths and angles for cell-cell contacts (*L*_*l*_ and *𝜙*). **B**. In the continuous limit, both the apical and basal caps are circular arcs with generic radius of curvature *ρ*, base *b* and contact angle *θ*. **C**. Diagram showing the relation between angles *θ*_*a*_, *θ*_*b*_ and *𝜙* at 3-fold vertices.

#### A.5 Existence of a symmetrical single-layered equilibrium state

As seen in our analysis of a tissue composed of wedge-shaped cells, for fixed 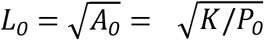, a bifurcation in the energy minima happens as parameter *T*_*a*_/*T*_*0*_ increases, with extrusion (*b*_*b*_ = *0* or *b*_*b*_ = *2b*) being favored for low *T*_*a*_/*T*_*0*_ and a symmetrical tissue configuration in which *b*_*b*_ = *b*_*a*_ = *b* becoming stable when *T*_*a*_/*T*_*0*_ is large. In fact, this equilibrium state exists for all values of *T*_*a*_/*T*_*0*_ > *0*.*5*, indicating that this system undergoes a supercritical pitchfork bifurcation.

As this is a special case of a tissue composed of wedge-shaped cells, we search for solutions of *∇E*_*cell*_ = *0*. Because of the tissue’s symmetry, *E*(*b*_*b*_, *h*) = *E*(*2b* − *b*_*b*_, *h*), which means *∂E*_*cell*_ /*∂b*_*b*_ = *0* when *b*_*b*_ = *b*_*a*_ = *b*. In that case, 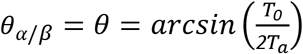, which necessitates *T*_*a*_/*T*_*0*_ > *0*.*5*. Under these conditions, equations (A18-19) simplify to

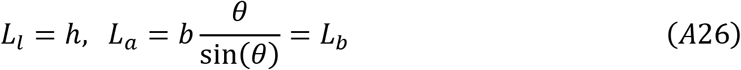

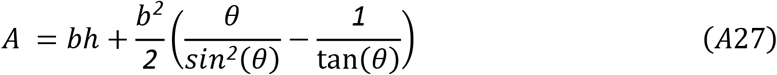

which depend only on variable *h*. Differentiating *E*_*cell*_ (Eq. A15) with respect to *h* gives us

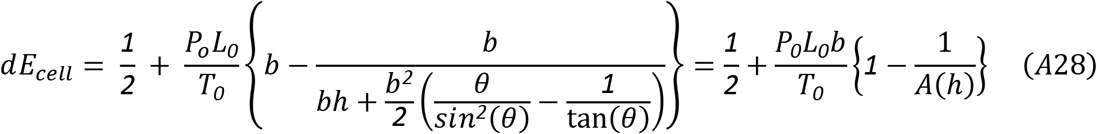

which, when equal to

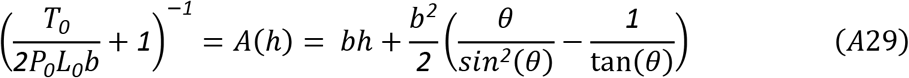

from which *h* may be obtained for any positive value of *b, P*_*0*_ and *L*_*0*_ as long as *T*_*a*_/*T*_*0*_ > *0*.*5*.

